# A systematic global review of mammalian carnivore responses to production forests

**DOI:** 10.1101/2023.03.26.534289

**Authors:** Evie M Jones, Amelia J Koch, Rodrigo K Hamede, Menna E Jones

## Abstract

1. Unmodified forests are increasingly rare worldwide, with forestry a major contributor to habitat modification. Extending conservation practices beyond protected areas is important to conserve forest ecosystems.
2. We investigate the response of native mammalian carnivores (both Order Carnivora and Family Dasyuridae) to production forests globally, including harvested native forest and timber plantations. We examine how carnivores recorded in production forests use these forests versus other land uses, particularly native and unharvested forest; how habitat use relates to threatened status, body size, diet, and harvesting method; carnivore responses to habitat features within production forests; and carnivore denning, breeding, and predation behaviour in production forests.
3. We review 294 studies recording 132 carnivore species in production forests. Carnivores generally show higher use of native and unharvested forests and lower use of agricultural land than production forests. Threatened species and large carnivores respond more negatively to production forests than non-threatened species and small carnivores respectively. Hypercarnivores respond more negatively than omnivores to plantations compared to native forest, but there was no difference in the use of harvested and unharvested native forest between these dietary groups.
4. Surprisingly, a high proportion of carnivore species use clearfelled more than unharvested native forest. In forest with partial harvesting or reduced-impact logging, most species show no difference in use between harvested and unharvested forest.
5. Carnivores generally respond positively to habitat features such as riparian areas and coarse woody debris. Several carnivores were recorded denning and breeding in production forests. Production forests often influence the prey availability, hunting success, and diet of carnivores.
6. We show that many carnivores use production forests, and how they respond to production forestry varies with species traits. We recommend that production forests are managed as valuable carnivore habitat, and highlight strategies to enhance the use of these forests by carnivores.

## Introduction

More than three quarters of the world’s land area has been modified by humans, and the loss of intact wilderness areas continues (Watson et al. 2016). The area of forests is declining worldwide, with 2.3 million square kilometres of forest lost between 2000 to 2012 (Hansen et al. 2013). While only 27% of this forest loss was attributed to permanent deforestation due to land use change (as opposed to temporary forest loss), the rate of deforestation is not diminishing (Curtis et al. 2018). Intact and protected habitats are no longer sufficient to preserve biodiversity and ecosystems (Hayes & Ostrom 2005).

Of all global forest loss between 2001 and 2015, 26% was due to forestry (Curtis et al. 2018). However, unlike commodity driven deforestation which involves the permanent loss of forest for other land uses, areas subject to forestry are regenerated so they can be harvested again (Curtis et al. 2018). Production forests, including harvested native forest and timber plantations, often retain much of their biodiversity and ecosystem functions (Edwards et al. 2014), particularly compared to other land uses such as agriculture (Brockerhoff et al. 2008). Production forests thus have the potential to provide valuable habitat for forest-dependent wildlife.

Mammalian carnivores – both placental (Order Carnivora) and marsupial (Family Dasyuridae) – are often sensitive to habitat degradation because they generally have large home ranges and high energy requirements, and their predatory nature brings them into conflict with humans (Carbone et al. 1999, Cardillo et al. 2004, Kosydar 2014, Marneweck et al. 2021). Loss of carnivores globally has cascading effects on ecosystems (Ripple et al. 2014), as carnivores play key roles in regulating prey (Ripple & Beschta 2012) and smaller predators (Crooks & Soule 1999), and in carrion recycling (Cunningham et al. 2018). These functions are often suppressed in modified landscapes, even where carnivores are present, due to low carnivore densities (Kuijper et al. 2016). The response of carnivore species to habitat modification varies, with some highly sensitive to and others able to adapt to disturbance (Crooks 2002). Recent research found 82% of large carnivore ranges globally fall outside Protected Areas (Braczkowski et al. 2023). Finding ways to improve carnivore conservation in modified landscapes is important for preserving carnivore populations and maintaining ecosystem function, and may even benefit landowners where carnivores can play a useful role in controlling populations of herbivores that damage trees and crops (Kuijper 2011, Davoli et al. 2022).

Ferreira et al. (2018) reviewed the literature on mammalian carnivore use of agroecosystems, including timber plantations which contained the highest number of species of all the agricultural habitats considered. They found non-threatened carnivores were more likely to use agroecosystems than threatened species, and that body size, trophic level and locomotion mode influenced carnivore use of these landscapes. However, their review only considered which species were recorded using plantations and did not compare use of plantations with unmodified habitats, or consider which habitat features in production forests influence carnivores (Ferreira et al. 2018). Understanding what landscape attributes influence carnivores to select or avoid production forests, including plantations, can help design management strategies that enhance the quality of these habitats for carnivores. To our knowledge, the global literature on carnivore use of harvested native forests has not been reviewed.

We investigate which native mammalian carnivore species use production forests worldwide, and how these species respond to these modified landscapes. Species for which meat contributes a substantial portion of their diet, i.e. dietary carnivores (including hypercarnivores and omnivores), occupy a distinct niche and face unique challenges relating to the need to find prey (Wolf & Ripple 2016, Nisi et al. 2022). Due to this, we define ‘carnivores’ as members of the placental Order Carnivora that are not strictly herbivorous (i.e. we exclude pandas [*Ailuropoda melanoleuca* and *Ailurus fulgens*] and some genet species), as well as the larger species (> 500 g) of marsupial carnivores in the Family Dasyuridae: Tasmanian devils (*Sarcophilus harrisii*) and quolls (*Dasyurus spp.*). These marsupial species occupy equivalent ecomorphological niches to different Families of placental carnivores (Jones et al. 2003, Glen & Dickman 2008) but tend to be overlooked in the literature. We review and summarise the literature reporting carnivores using production forests, separating studies into two broad categories: plantations and harvested native forest. We synthesise the information to discover what factors influence directional responses (positive or negative) of carnivores to production forests compared to other habitat types, and which habitat features affect carnivore use of these forests. This information could be used to develop management practices that improve the value of these landscapes to carnivores. We aim to answer the following questions:

1. What is the geographic distribution of global studies of carnivores in production forests, and what is the focus of these publications?
2. How do carnivores use production forests versus intact habitat and other land uses, and how is habitat use influenced by threatened status, diet and body size?
3. How does carnivore use of harvested native forests vary with time since harvesting and harvesting method?
4. What habitat features influence carnivore use of production forests?
5. How do production forests influence carnivore predation, denning and breeding behaviour?
6. What are the knowledge gaps in relation to carnivores in production forests?

## Methods

### Search strategy

We conducted a systematic search for published papers, unpublished theses and reports up to January 2023 on the topics of mammalian carnivores and production forests. We defined carnivores as any species from the placental Order Carnivora, excluding strict herbivores and frugivores, and marsupial carnivores in the Family Dasyuridae larger than 500 g because most smaller dasyurids are insectivorous rather than carnivorous (Berkovitz & Shellis 2018). We did not place a weight limit on placental carnivores as we found few species < 500 g and those are mostly small mustelids, which are carnivorous. Production forest was defined as land managed primarily for wood and fibre production. We reviewed publications from the databases SCOPUS (www.scopus.com) and Web of Science (webofscience.com) as well as theses from Open Access Theses and Dissertations (OATD: oatd.org) using the terms presented in Table 1. We designed the search strings as follows: [any terms in Concept 1] AND [any terms in Concept 2] AND NOT [any terms in Exclusions]. Search terms were chosen to identify community-wide studies of mammals as well as those centred only on carnivores.

**Table 1:**
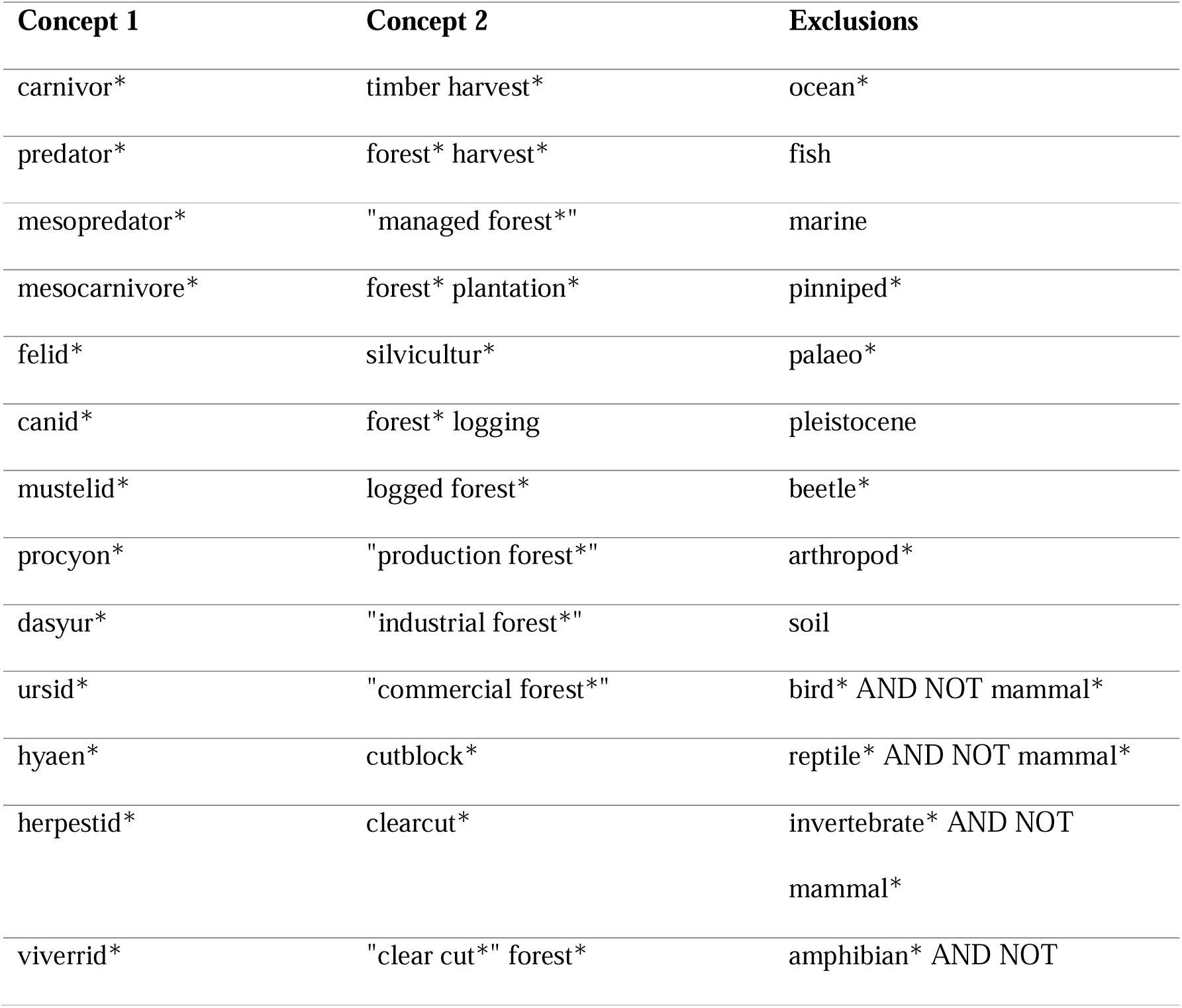
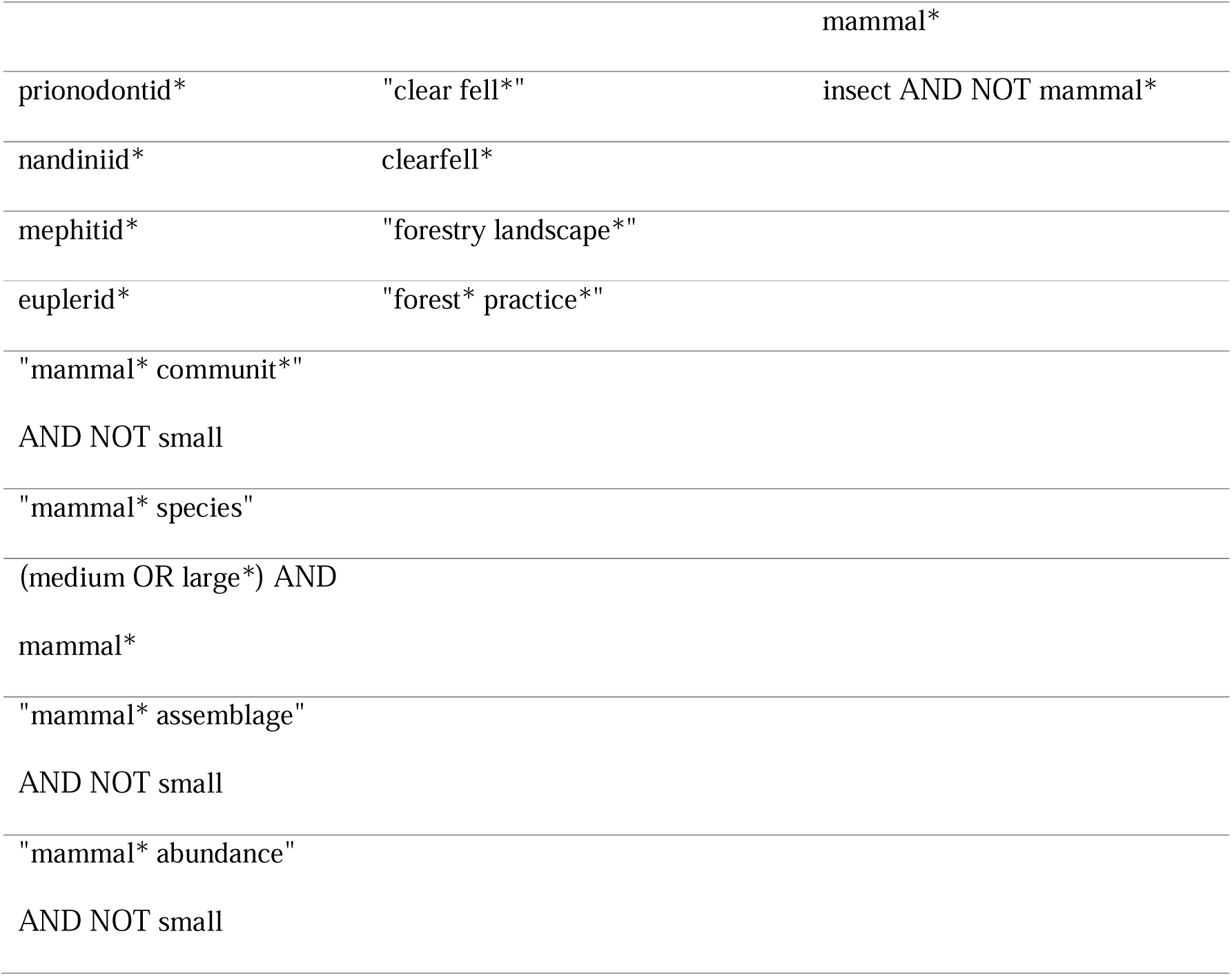
Search terms used to identify relevant publications from literature databases.

The search metrics were recorded for each set of terms at each stage of the search. Abstracts from the full search were uploaded into the screening and data extraction software Covidence (covidence.org) (Veritas Health Innovation 2022), which is designed specifically to support systematic literature reviews. Two authors (first and last) screened all abstracts and full texts, conferring and coming to a consensus where decisions on whether to accept or reject were conflicted. Publications discovered through this search were excluded if:

1. They did not report any carnivore species using production forests;
2. They were a review of other literature, or involved modelling only, and did not include primary empirical observational or experimental research;
3. We could not distinguish the use of production from intact forest by carnivores, such as when these were not considered separately in the study;
4. The study was not published in English;
5. The study included only carnivores that were not native to the region.

After reviewing the publications and applying filters, 294 published and unpublished studies remained.

### Data compilation

For all studies we recorded the country or region in which the study was located, the survey method, the response variable considered, and all carnivore species found in the production forest. We used the IUCN Red List of Endangered Species to classify each carnivore species as ‘non-threatened’ (Least Concern) or ‘threatened’ (Vulnerable, Endangered, or Critically Endangered). We identified the body size and diet (hypercarnivore – where the vast majority of the species’ diet is comprised of meat – or omnivore) of each species using the Encyclopedia of Life (https://eol.org/) and Animal Diversity Web (https://animaldiversity.org/). We divided carnivores into ‘small’ (< 21.5 kg) and ‘large’ (≥ 21.5 kg) based on the approximate body size at which predators switch from small to large prey (Carbone et al. 1999).

We divided the studies into two categories: those considering carnivore use of timber plantations (‘plantations’), and those considering carnivore use of harvested native forest. We separated these studies as the ecological value of plantations and harvested native forest differ: harvested native forest provides a more ‘natural’ forest habitat, while plantations involve more severe habitat modification, but can reduce the overall area harvested compared to harvested native forest (Brockerhoff et al. 2008). This means species often respond differently to the two land uses (Chaudhary et al. 2016). Some studies were included in both categories. For the studies of harvested native forest we considered harvesting method, classifying these where possible into the most commonly reported methods: clearfelling, partial harvesting, and reduced-impact logging (RIL) (see Table 2 for definitions).

**Table 2:** Definitions of key terms used in the study.

| Term | Definition |
| --- | --- |
| Habitat use | Significant differences (or lack thereof) in abundance, density, occupancy, distribution, presence, or habitat selection of species. |
| Undergrowth | Includes understorey, ground cover, visual obstruction. |
| Coarse woody debris (CWD) | Includes logs, downed trees, logging debris, other woody debris. |
| Riparian areas | Includes rivers, lakes, wetlands. |
| Habitat edge | Edges between forest and non-forest, or plantation and native forest. |
| Stand age | Time since logging. As age classes differed among studies, we grouped them as follows: |
| - Younger | - The youngest age class considered in the study. |
| - Mid-stage regenerating | - Neither the youngest nor oldest age class considered in the study. |
| - Older/mature | - The oldest age class considered in the study. |
| Harvesting method | Forest harvesting type used in the study landscape (native forest only). |
| - Clearfelling | - Removal of all trees in the harvest footprint. |
| - Partial harvesting | - Removal of only some trees within the harvest footprint, including selective logging. |
| - Reduced impact logging (RIL) | - A logging method used in tropical countries, designed to minimise environmental damage while maximising efficiency (Holmes et al. 2002). |

To investigate patterns of habitat use by carnivores in production forest landscapes, we used papers with a response variable that could be considered ‘habitat use’, such as abundance, density, occupancy, distribution, presence, or habitat selection (Table 2). We summarised the number of species that showed differences (or no difference) in their use of different vegetation and human land-use types: plantation versus native forest (both harvested and unharvested, as many studies did not specify this), native grassland/scrub, or agricultural land; and harvested versus unharvested native forest. ‘Higher use’ was defined as a more positive response (e.g. higher abundance, positive selection) to one habitat compared to another that was reported as statistically significant (e.g. p < 0.05 or confidence intervals not overlapping zero), while no significant difference was defined as ‘no difference’ (Table 2). Some species were reported to have different responses by different studies, and these species were counted once for each reported response.

To investigate the influence of diet, threatened status and body size on habitat use by carnivore species in production forests, we summarised the number of species that were categorised as hypercarnivores versus omnivores, non-threatened versus threatened, and large vs small, that showed differences (or no difference) in their use of plantations versus native forest, and harvested versus unharvested native forest.

To identify the influence of native forest harvesting on carnivores, we compared carnivore use of harvested versus unharvested forest among the different harvesting methods. We also investigated the responses of carnivores to different stand ages (i.e. time since logging), grouping stand ages into younger, mid-stage regenerating, and older/mature age classes (Table 2).

To examine how carnivores respond to different habitat features in production forests, we identified statistically significant responses (‘positive’ or ‘negative’) by carnivore species to any reported habitat features in production forests and looked for commonly reported features (e.g. undergrowth, riparian areas, coarse woody debris [CWD] [Table 2]). We also considered responses of carnivores to forestry roads and habitat edges (Table 2). For all species reported denning in production forests, we identified common den site features. We also summarised observations on diet, prey availability, hunting behaviour/success, and reproductive success of carnivores in production forests.

## Results

### Focus of publications

A total of 294 papers from 44 countries reported carnivores using production forests: 34% from North America, 28% from Asia, 16% from Europe, 14% from Central/South America, 5% from Africa/Middle East, and 2% from Oceania (Figure 1, Appendix S1, S2). North America had by far the most studies on carnivores in harvested native forest, followed by Asia, while studies focussing on carnivores in plantations were conducted mainly in South America, Europe and Asia (Figure 1). The vast majority of studies focussed on habitat use (98 in plantations, 171 in harvested native forest), while 11 considered diet (six in plantation, five in harvested native forest), and 29 considered hunting success/prey availability (six in plantation, 23 in harvested native forest). Several other topics were only studied in harvested native forest: seven studies on reproductive success, one on mortality, one on extinction risk, one on gene flow, and one on health (stress/body condition). Some studies considered multiple topics.

**Figure 1:**
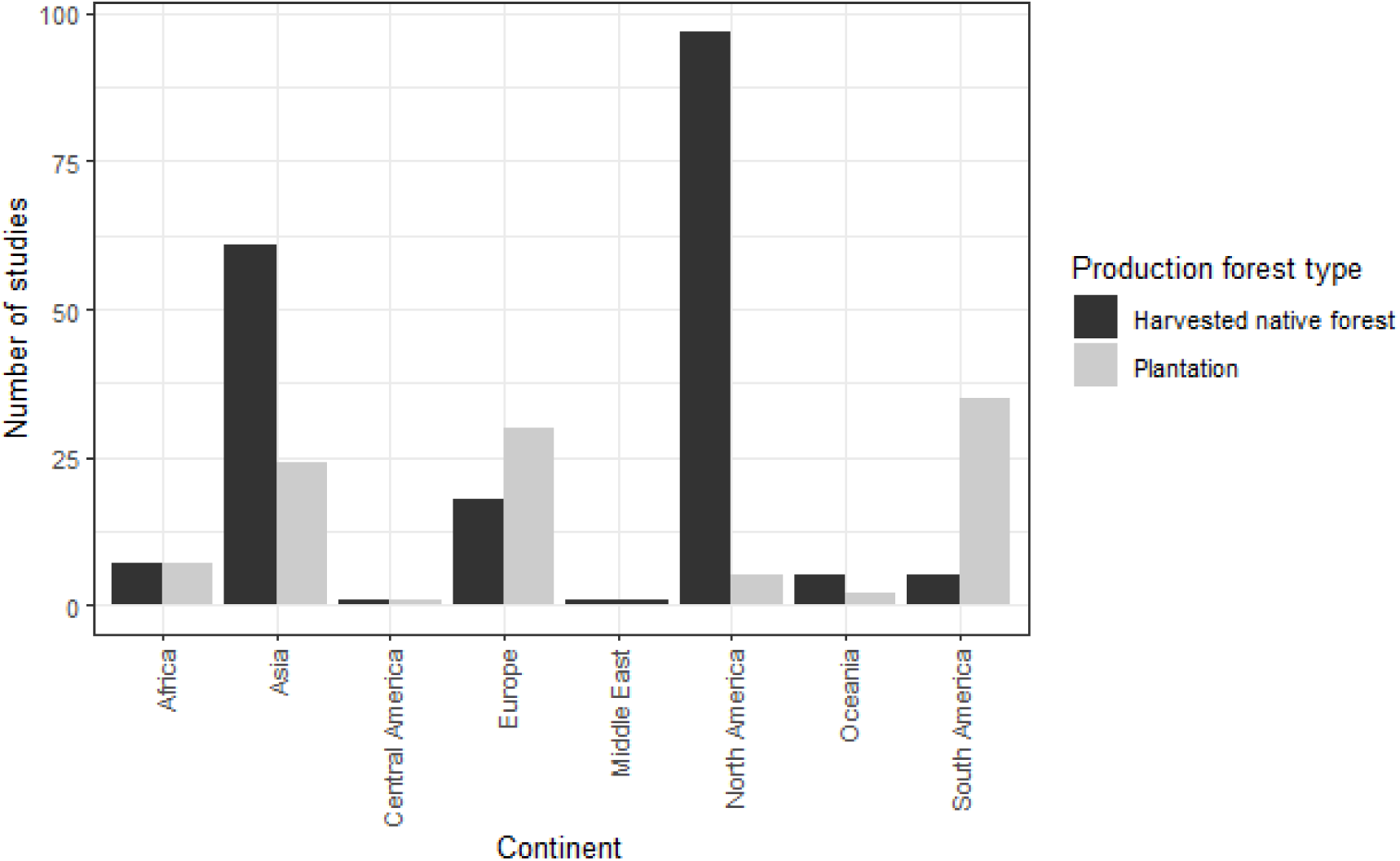
Number of studies conducted in each continent that reported native mammalian carnivores using production forests.

### Carnivores reported in production forests

A total of 132 carnivore species were reported using production forests: 91 in plantations, 90 in harvested native forest, and 49 in both (Appendix S3). Of the 291 Carnivora species recorded on the IUCN red list that were not ‘Extinct’, 45% were recorded in production forests (this is an underestimate as our methods excluded strict herbivores in the Order Carnivora), along with four of the seven Dasyuridae species considered.

All terrestrial Carnivora families were represented (excepting Ailuridae, which was excluded from our analysis due to containing only the herbivorous red panda), with the addition of the marsupial carnivore Family Dasyuridae (Figure 2). Species in the placental families Nandiniidae, Prionodontidae and Eupleridae were not reported in plantations, and Hyaenidae were reported only in plantations, while all other families occurred in both plantations and harvested native forest. The Family Eupleridae is endemic to Madagascar, and we did not find any studies in plantations in that country, explaining the lack of records of these species in plantations. Mustelidae and Felidae were the two most commonly reported families in both harvested native forest and plantations, closely followed by Canidae in plantations. For most carnivore families, there were similar numbers of species recorded in plantation as in harvested native forest (Figure 2). One notable exception to this is Canidae, which had more than twice the number of species recorded in plantations than in harvested native forest.

**Figure 2:**
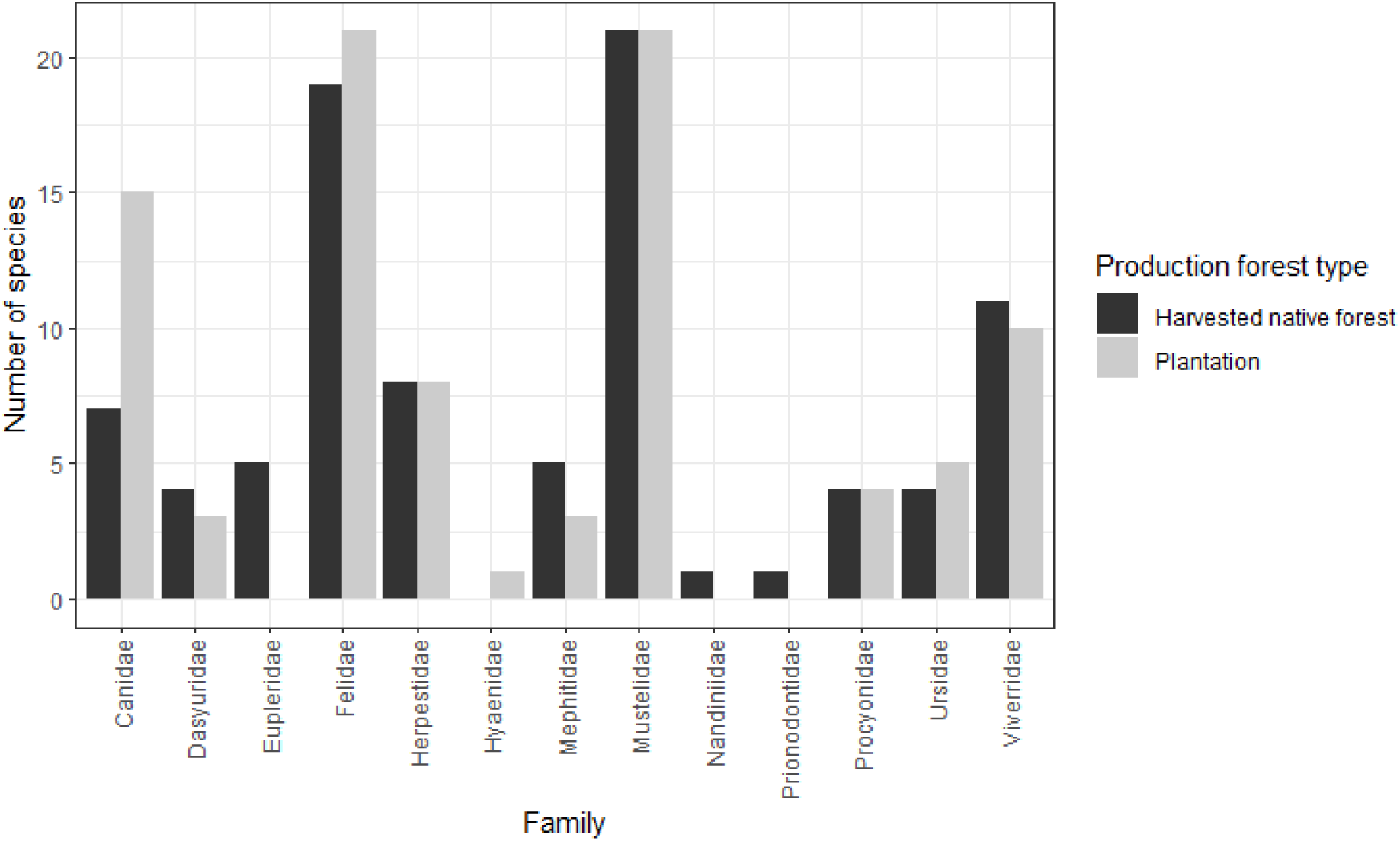
Number of species in each carnivore Family reported using production forests.

### Carnivore use of plantations and harvested native forest compared to other habitats

Carnivores generally used plantations less than native forest habitats and more than agricultural land, with no difference in use between plantations and native grassland/scrub (Figure 3). Where other land uses were considered alongside harvested native forest, carnivores tended to respond negatively to agricultural land (including oil palm) (eight species, seven studies) and human settlements (six species, seven studies), although a positive response to agricultural land and human settlements was found for three and two species respectively (Appendix S5).

**Figure 3:**
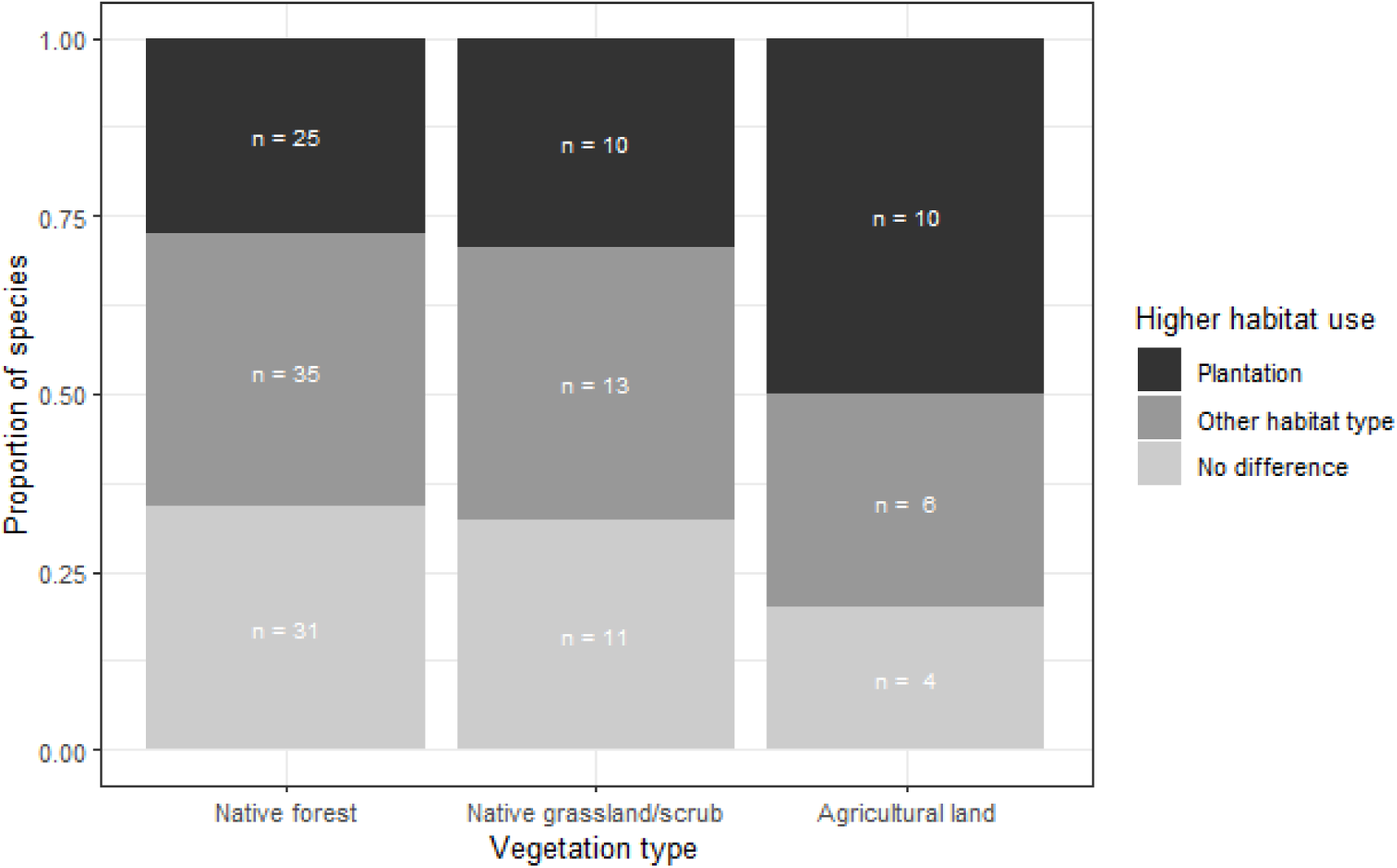
Carnivore use of plantations compared to other habitat types.

Species traits of carnivores appeared to influence their habitat use in production forests. The highest proportion of hypercarnivores appeared to show greater use of native forest than plantations, with only a few species showing the reverse (Figure 4a). Omnivores also seemed to respond more positively to native forest than plantations, but the difference was much less than that shown by hypercarnivores. Both hypercarnivores and omnivores tended to show no difference in use between harvested and unharvested native forest.

**Figure 4:**
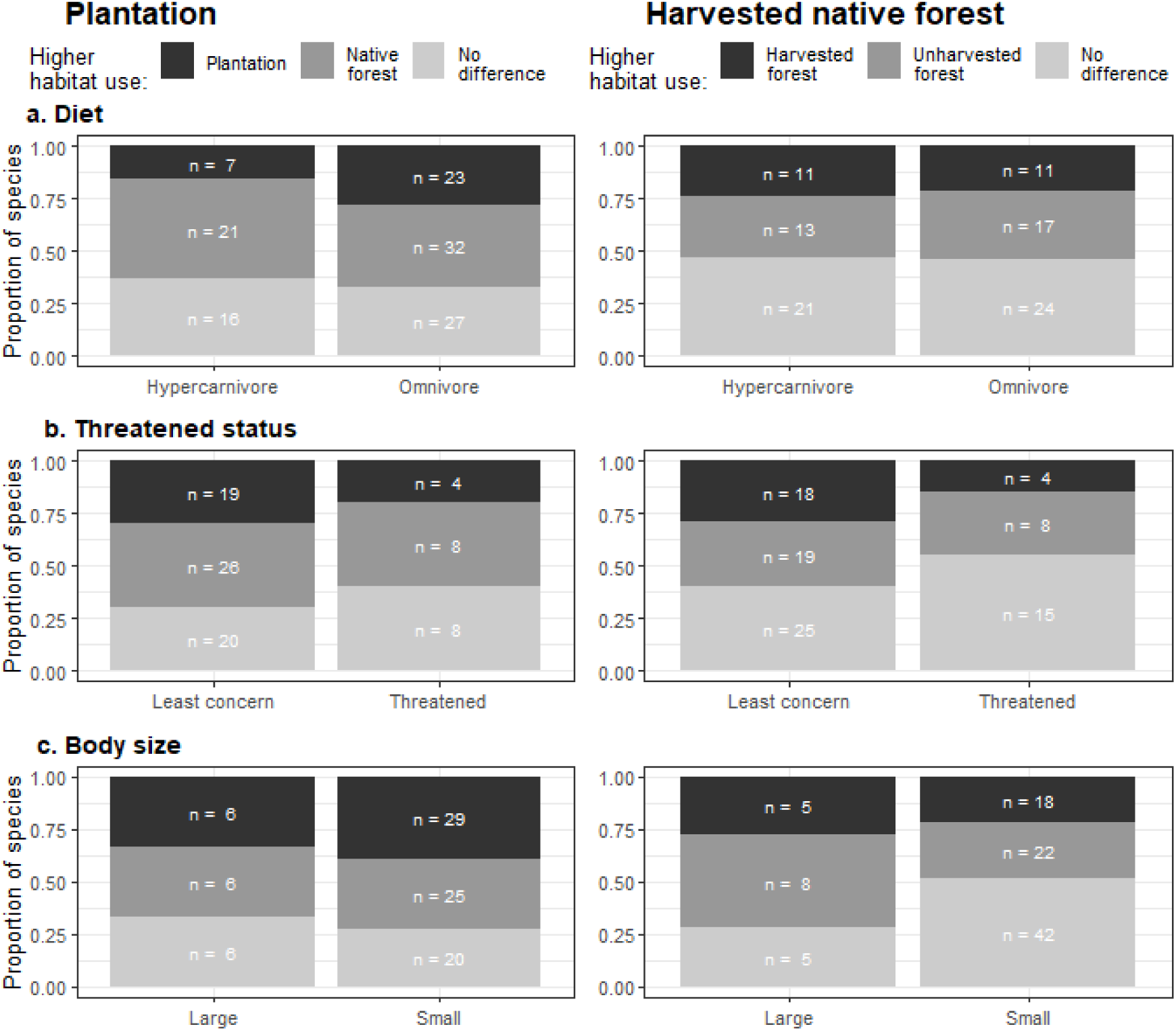
Statistically significant differences in habitat use of plantations versus native forest, and harvested versus unharvested native forest, by carnivore species in relation to diet, IUCN threatened status and body size (small < 21.5 kg, large ≥ 21.5 kg).

Threatened species appeared to respond more negatively to production forests than non-threatened species. Non-threatened species were more likely to show higher use of production forests compared to native/unharvested forest (Figure 4b). A smaller proportion of all existing threatened species (21%) than non-threatened species (34%) were reported to occur in plantations; while in harvested native forest the proportions of all threatened and non-threatened species reported were similar (31% and 30% respectively). Note these are underestimates as we excluded strict herbivores from our analysis, but we were comparing them to the total number of threatened/non-threatened species in Carnivora, which include some herbivores.

Large carnivores responded more negatively to production forests than small carnivores. A higher proportion of large than small carnivores used unharvested more than harvested native forest, and a higher proportion of small than large carnivores used plantations more than native forest (Figure 4c). Small carnivores made up over 80% of all carnivores using production forests.

### Responses of carnivores to forest harvesting

In native forest, carnivores showed different responses to different forms of harvesting. Unexpectedly, in native forest where clearfelling was used as a harvesting method, more than a third of carnivores used harvested more than unharvested forest (Figure 5). In comparison, when the forest harvesting method was partial harvesting or RIL, nearly half of species and nearly all species respectively showed no difference in use of harvested versus unharvested forest.

**Figure 5:**
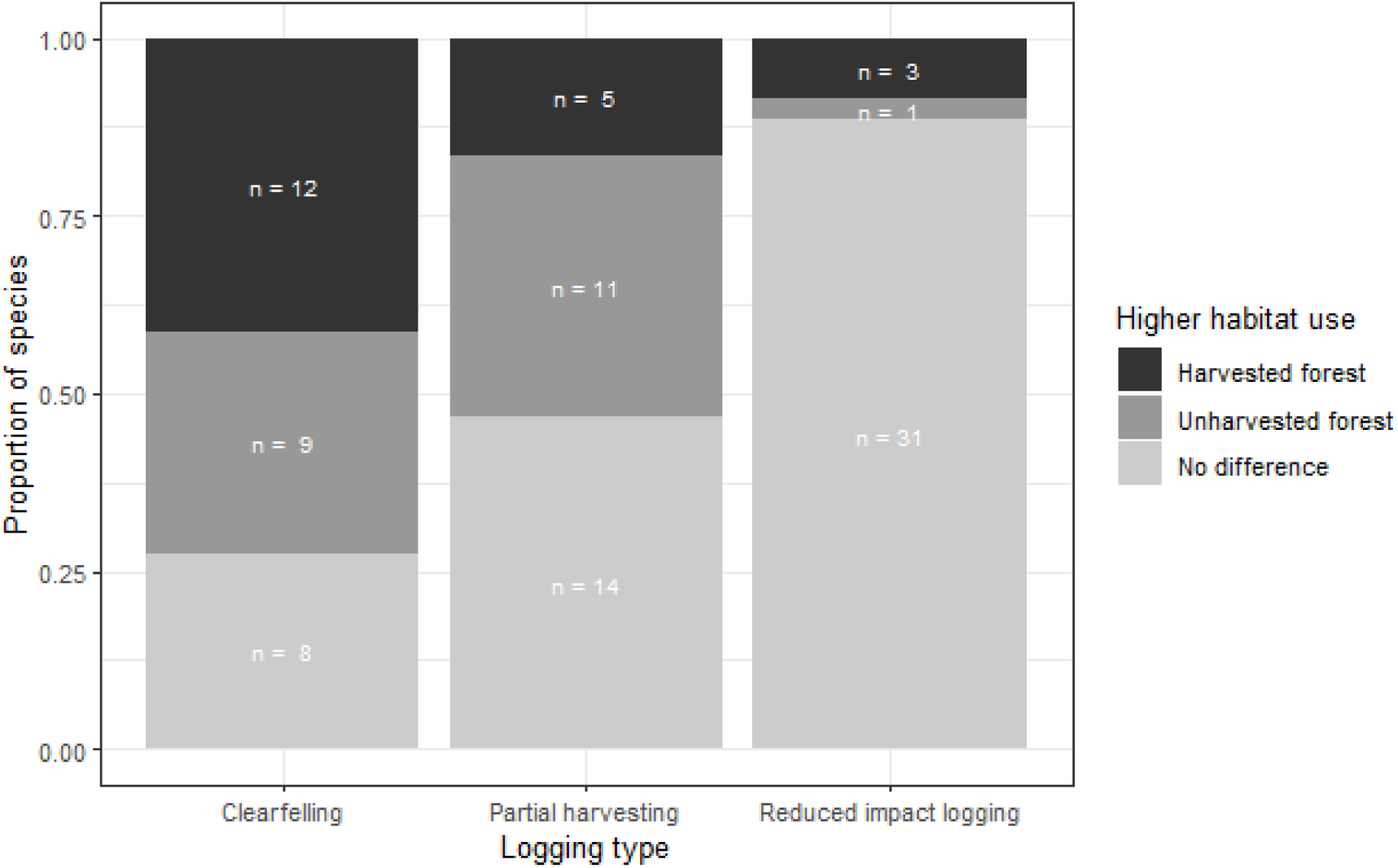
Statistically significant differences in habitat use of harvested versus unharvested forest by carnivore species in native forests with different harvesting methods.

The response of carnivores to harvesting-related habitat features varied. Carnivores responded positively and negatively to recent clearcuts in production forests, with some species recorded with different responses by different studies (Table 3). In harvested native forest, selection among stand ages also differed among carnivores (Table 3). There was limited information on the response of carnivores to stand ages in plantations.

**Table 3:** Responses of carnivore species to habitat features in production forests, including the number of studies recording these results. See Appendix S4 for specific studies.

| Habitat feature | Plantations |  | Harvested native forest |  |
| --- | --- | --- | --- | --- |
|  | *Positive response* | *Negative response* | *Positive response* | *Negative response* |
| Stand age |  |  |  |  |
| - Younger |  |  | 12 species<br>12 studies |  |
| - Mid-stage regenerating |  |  | 5 species<br>7 studies |  |
| - Older/mature |  |  | 8 species<br>15 studies |  |
| Clearcuts | 5 species<br>4 studies |  | 4 species<br>5 studies | 6 species<br>9 studies |
| Undergrowth | 11 species<br>14 studies | 3 species<br>3 studies | 5 species<br>7 studies | 5 species<br>5 studies |
| Canopy cover | 3 species<br>3 studies | 3 species<br>3 studies | 8 species<br>14 studies | 5 species<br>6 studies |
| Coarse woody debris (CWD) | 4 species<br>5 studies |  | 6 species<br>7 studies |  |
| Larger/taller trees |  |  | 6 species<br>7 studies |  |
| Riparian areas | 9 species<br>8 studies |  | 8 species<br>15 studies |  |
| Forestry roads | 3 species<br>3 studies | 3 species<br>3 studies | 11 species<br>14 studies | 13 species<br>14 studies |
| Habitat edges | 7 species<br>2 studies | 1 species<br>1 study | 6 species<br>9 studies | 4 species<br>5 studies |
| Den features |  |  |  |  |
| - Large trees |  |  | 4 species<br>4 studies |  |
| - Logs |  |  | 3 species<br>6 studies |  |
| - Tree cavities |  |  | 3 species<br>5 studies |  |
| - Dead trees |  |  | 3 species<br>3 studies |  |
| - Logging debris |  |  | 5 species<br>7 studies |  |

### Habitat features important to carnivores in production forests

Some habitat features were commonly reported as important for carnivores in production forests. In plantations, carnivores tended to respond positively to more undergrowth. In harvested native forest, the response to undergrowth was less consistent, with equal numbers of species responding positively to more and less undergrowth (Table 3). Response to canopy cover was variable in both plantation and harvested native forest (Table 3). CWD generally positively influenced carnivores in plantations and harvested native forest (Table 3). Larger/taller trees positively influenced six species in harvested native forest (Table 3). Riparian areas were important for many carnivores in plantations and harvested native forest (Table 3).

The response of carnivores to forestry roads was likewise highly variable, with similar numbers of species responding positively and negatively to roads (Table 3), and many showing no response (20 species, six studies, Appendix S5). Carnivores sometimes responded positively and sometimes negatively to habitat edges (Table 3).

### Diet, denning and breeding of carnivores in production forests

Carnivore denning behaviour was considered in 20 studies in harvested native forest and seven in plantations. Due to the limited number of dens found in plantations, we only considered the harvested native forest studies. Six of the denning studies were on American marten (*Martes americana*). Five species were found with dens in logging debris, including maternal dens of two species: American black bears (*Ursus americana*) (White et al. 2001) and wolverines (*Gulo gulo*) (Scrafford et al. 2017). Other important denning features were large trees, logs, tree cavities (three species, five studies), and dead trees (Table 3). Seven studies (five species) considered reproductive success of carnivores (Appendix S5), all reporting successful breeding in production forests, although three found lower success in harvested than mature forest (White et al. 2001, Kosterman et al. 2018, Holbrook et al. 2019).

The diet and predation success of many carnivores was influenced by production forests. Some studies reported higher hunting success (Bojarska et al. 2017, Ausilio et al. 2022) or higher prey availability (Soutiere 1979, Boisjoly et al. 2010, Parsons et al. 2020) for carnivores in harvested than unharvested native forest, which sometimes caused them to select this habitat (Simons-Legaard et al. 2016, Gagné et al. 2016, Roffler et al. 2018, Olson et al. 2023) but not always (Hargis et al. 1999). In other cases prey availability or hunting success was lower in harvested than unharvested native forest (Andruskiw et al. 2008, Rayan & Linkie 2015, Jiang et al. 2017). Some species changed their prey selection between plantations and native forest (Kaneko et al. 2009, Moreira-Arce et al. 2015, Twining et al. 2019) and between harvested and unharvested forest (Sidorovich et al. 2010, Gervasi et al. 2013).

## Discussion

Our global review highlights the wide variety of native mammalian carnivore species that use production forests, reinforcing the need to manage these landscapes as valuable carnivore habitat. We show that many carnivore species can successfully hunt, den, and breed in production forests. Carnivores vary in their response to production forests, with some showing higher use of production forests than intact forests, some the reverse, and many no difference in use between habitats. Species traits influence the responses of carnivores to plantations, with hypercarnivores, large carnivores and threatened species responding more negatively to plantations than omnivores, small carnivores and non-threatened species, although these traits were less important in predicting carnivore responses to harvested native forest.

A couple of caveats are important. First, we assessed the comparative responses of carnivores to production forests and other habitats in relation to their ‘habitat use’, which in our study was broadly defined and included measures such as occupancy, abundance, and habitat selection (Table 2). This allowed us to make some broad generalisations about the use of different habitats by carnivores, but the results should be interpreted with caution as they include multiple different response variables considered together, e.g. ‘higher use’ by carnivores could mean higher abundance, greater occupancy, or selection for that habitat. Additionally, our search terms precluded reporting of species that may avoid production forests entirely. Some carnivore species that are highly sensitive to habitat fragmentation and disturbance may rarely or never be found in production forests and require intact habitat to survive (Ferreira et al. 2018). Our study shows only the habitat use of carnivores that do use production forests.

The majority of research on carnivores in production forests has been undertaken in North America and Asia, with few studies from Africa and Oceania. Asia contains more plantation forests than anywhere else in the world (FAO 2020), as well as extensive tropical forest harvesting (Edwards et al. 2014) and high species richness of carnivores (Marneweck et al. 2021) explaining the high number of studies conducted there. North America likewise contains species-rich carnivore communities (Marneweck et al. 2021). Most of the research there was focussed on harvested native forest rather than plantations, and Canada and the USA are forest-rich countries, ranking third and fifth respectively in the list of countries with the most primary forest (FAO 2020). The focus on North America likely reflects this extensive forest cover as well as the importance of the forestry industry in these countries. The limited research in Oceania can be explained by only eight native terrestrial mammalian carnivore species > 500 g present there, all of which are in the Family Dasyuridae, except the dingo (*Canis dingo*). Carnivores in production forests in Africa remain understudied, despite the continent harbouring about a third of all carnivore species (Harris et al. 2022) and 16% of the world’s forests. 60% of all Africa’s forest is not primary forest, indicating forestry is a major land use, though the continent contains only 2% of the world’s plantations (FAO 2020).

Interestingly, most carnivore families had similar numbers of species recorded in plantations and harvested native forest. This is contrary to expectations as biodiversity tends to be lower in plantations than harvested native forest (Chaudhary et al. 2016), and is unlikely to be due to sampling bias as two-thirds of the papers in this review were conducted in harvested native forest. This may indicate that carnivore families generally have a similar likelihood of presence in both production forest types, although we cannot say if this is the case due to few studies that explicitly compared the use of plantations to harvested native forest by carnivores. Notably, as discussed below, a species’ presence in a forest type does not show the value of the habitat to the animal, such as to what degree they use that habitat and which elements are important (Johnson 1980).

There were substantially more canid species recorded in plantation than harvested native forest. The Canidae Family contains many generalist and adaptable species, and has diversified to occupy a wide range of habitat types, making it one of the most geographically widespread carnivore families (Padilla & Hilton 2015). This may allow them to better adapt to human-modified landscapes such as plantations than other carnivore families. For instance, the culpeo (*Lycalopex culpaeus*), a generalist canid that thrives in a variety of habitats (Malo et al. 2021), was frequently found to use plantations more than native forest (Acosta-Jamett & Simonetti 2004, Guerrero et al. 2006, Escudero-Páez et al. 2018).

### Influence of species traits on carnivore responses to production forests

Diet and threatened status appeared to influence the response of carnivores to plantations, but less so for harvested native forest. Hypercarnivores tended to use native forest more than plantations. These results are consistent with Ferreira et al. (2018) who found that most carnivore species using agroecosystems were generalist consumers. Generalist species often tolerate human-modified landscapes better than specialists (Law et al. 2017), and often even benefit from these landscapes, examples being the crab-eating fox (*Cerdocyon thous*) (Coelho et al. 2014) and maned wolf (*Chrysocyon brachyurus*) (Lyra-Jorge et al. 2008). Similarly, threatened species generally show a less positive response to production forests than non-threatened species, and a lower proportion of total threatened species than non-threatened species were found in plantations, indicating threatened species are more likely to be absent from plantations. This is also supported by Ferreira et al. (2018) who found threatened species were much less likely to use agroecosystems than non-threatened species. These results are unsurprising as habitat loss and fragmentation is a major cause of global carnivore declines (Ripple et al. 2014).

Within harvested native forest landscapes, there was no major difference between the responses of hypercarnivores and omnivores to harvested versus unharvested native forest, and the proportions of all threatened and non-threatened species reported were similar. Native forest harvesting has a lower impact on biodiversity than plantations (Chaudhary et al. 2016), and our results indicate that carnivores likewise respond more negatively to plantations than harvested native forest, likely due to harvested native forest retaining more of the original habitat features than plantations. This is despite similar numbers of carnivore species recorded overall in plantations and harvested native forest, which reinforces the value of comparing the degree of habitat use among different land types, allowing detection of patterns that are not revealed with simple presence/absence analyses. As stated above, the actual proportion of carnivores that respond negatively to production forests is likely to be higher than our study shows, as we excluded carnivores that would only use intact native forests.

Large carnivores generally show higher use of unharvested than harvested native forest. Similarly, small carnivores show a slightly more positive response to plantations than do large carnivores. Large carnivores are vulnerable to landscape modification due to conflict with humans, high energy requirements, and large home ranges (Carbone et al. 1999, Cardillo et al. 2004, Kosydar 2014). In particular, large carnivores are reliant on large prey, which may be better provided for in intact landscapes (Carbone et al. 1999).

### Responses of carnivores to production forest landscape features

Forest harvesting is generally perceived as detrimental to wildlife, and clearfelling – which involves removing all vegetation within the harvest footprint – is more detrimental to species richness than other harvest types (Chaudhary et al. 2016). Surprisingly, a higher proportion of carnivores responded positively to clearfelling than other harvesting methods in native forest. Likewise, while several carnivore species responded negatively to recent clearcuts, others responded positively to them. This is likely due to increased prey availability or hunting success in cleared areas (Gagné et al. 2016, Ausilio et al. 2022). Carnivores in forest harvested using reduced impact logging (RIL), or to a lesser extent partial harvesting, generally showed no difference in use between harvested and unharvested forest. These methods do not involve removal of all trees in the harvest footprint, allowing some habitat and structural characteristics to persist (Man et al. 2008). For many carnivores, the habitat changes caused by these methods may not be severe enough to cause changes in their use of these habitats. This response was particularly marked with RIL, where few species were recorded with a difference in use between harvested and unharvested forest. RIL is often implemented in tropical countries such as Brazil (Carvalho Jr et al. 2021) and Indonesia (Jati et al. 2018) due to concern about the sustainability of harvesting these tropical forests, and is designed to minimise impacts on the environment while maximising output (Holmes et al. 2002). RIL is often more profitable than conventional logging (Holmes 2015) and less damaging to biodiversity than other harvesting methods (Bicknell et al. 2014, Chaudhary et al. 2016). Our results suggest RIL is likely to be successful at preserving carnivore populations in logged tropical forests, consistent with other research that has found little response of large mammals to RIL (Bicknell et al. 2015).

One challenge common to all carnivores in our study is the need to find prey, which is often influenced by production forest landscapes. Some studies report higher prey availability or hunting success in harvested than unharvested native forest, others lower. One study reported grey wolves (*Canis lupus*) using forestry fences to help kill red deer (*Cervus elaphus*) (Bojarska et al. 2017). Similarly, wolves take advantage of forestry roads for ease of movement, which increases mortality of threatened caribou (*Rangifer tarandus caribou*) (Vanlandeghem et al. 2021). These examples demonstrate how the influence of production forests on carnivores, even when positive, can have knock-on effects to other parts of the ecosystem. While some carnivores take advantage of roads and edges, overall carnivore responses to forestry roads and edges are variable. Roads and edges can provide hunting, travel and scavenging opportunities for predators (Gurarie et al. 2011, Bojarska et al. 2017, Andersen et al. 2017), but may also expose them to anthropogenic threats (Thiel 1985, Jones 2000, Balme et al. 2010), explaining why some species select for these features and others avoid them.

Carnivores generally respond more positively to production forests than agricultural land and human settlements, suggesting production forests may provide more valuable habitat to carnivores than other modified landscapes. Ferreira et al. (2018) similarly found that plantations supported more carnivore species than agricultural land. Production forests usually contain more structural diversity than agricultural land, such as understorey vegetation, which supports species rich mammal communities (Simonetti et al. 2013).

### Management implications and knowledge gaps

Our results suggest several management strategies could enhance the value of production forests to carnivores. We found riparian areas and native/unharvested forest remnants to be valuable to a wide range of carnivores, so maintaining a heterogeneous landscape with a mosaic of plantations/harvested forest along with a network of native/unharvested forest and riparian areas is likely to benefit carnivores and other species (Tews et al. 2004, Lindenmayer & Hobbs 2004). We also found different carnivore species used different stand ages of harvested native forest to varying extents, so maintaining a mosaic of forest ages may benefit carnivores by providing the forest age classes selected by different species and ensuring preferred stand ages are always available in the landscape. At a fine scale, maintaining diverse undergrowth (particularly in plantations) and abundant CWD is likely to improve the habitat quality of production forests for carnivores, as these habitat features were valuable to many species. Carnivores regularly denned in large trees, logs, tree cavities, and dead trees, so preserving these features may provide denning opportunities. Debris from timber harvesting was also found to supply valuable denning habitat for many species.

Managing production forests to enhance carnivore populations may benefit the forestry industry as well as carnivores themselves by allowing carnivores to control browsing prey populations. Browsing mammals cause significant damage to regenerating and planted saplings (Wallgren et al. 2013), and may be culled at substantial cost to producers (FPA 2017). In managed forests, the impacts of carnivores on browsers is often altered or absent, allowing herbivore numbers to increase (Kuijper 2011). Improving carnivore conservation in production forests may allow them to act as a biological pest control, as with crops in some instances (Williams et al. 2018, Davoli et al. 2022). If effective, this could save damage to trees and reduce the costs and ethical quandaries surrounding culling, and increase impetus to preserve carnivore populations in production forests.

While habitat use by carnivores in production forests has been well studied, some key knowledge gaps remain. Only one study considered carnivore stress and health in production forests, finding that stress in brown bears (*Ursus arctos*) following forest harvesting was lower for females and higher for males, and that bears in harvested areas had lower body condition (Bourbonnais 2013). Studying physiological responses to habitat change can help identify environmental stressors and their impact on species (Wikelski & Cooke 2006, Coristine et al. 2014). Logging and forest fragmentation are known to affect stress levels in other species (Suorsa et al. 2003, Leshyk et al. 2012), and chronic stress can impact reproduction (Sheriff et al. 2009, Narayan & Hero 2014), immunity (Owen et al. 2012) and survival (Bradley 1987), potentially causing population-level effects such as declines in abundance and reduced resilience to disease. Chronic stress in carnivores in production forests is therefore an important question to address. Similarly, only one study considered carnivore mortality in production forests, finding higher mortality of American martens in harvested than unharvested forest, both from natural and human causes (Thompson & Colgan 1994). Identifying the stress, health and population dynamics of carnivores in production forests and unmodified landscapes could help elucidate the responses of carnivores to these landscapes beyond habitat selection.

### Conclusions

We demonstrate that a wide range of carnivore species use production forests, and that their responses to these landscapes relate to species traits such as diet, body size and threatened status. Many carnivores, generally the smaller and more generalist species, were not only able to survive in production forests, but thrived in them, with a more positive response to production forests than unmodified forest. However, other species, particularly hypercarnivores, large carnivores and threatened species, had a more negative response to production forests, emphasising the importance of retaining unmodified forest to support these species. Management techniques to enhance the habitat quality of production forests may also increase the use of these landscapes by more sensitive carnivore species.

Habitat loss and modification is a major threat to carnivores worldwide, with cascading effects on ecosystems (Ripple et al. 2014). With 82% of large carnivore ranges falling outside of Protected Areas (Braczkowski et al. 2023), modified landscapes are increasingly important for carnivore conservation. We show that production forests can provide valuable habitat for many carnivores, and forest management techniques can improve habitat quality for these species. Production forests can make a valuable contribution to carnivore conservation globally if managed with this goal in mind.

