## Appendices for "A systematic global review of mammalian carnivore responses to production forests"

**Appendix S1:** Summary of papers reporting carnivores using timber plantations, including the country in which the research was conducted, the response variable considered, whether the response variable fit our definition of ‘habitat use’, the species recorded, and where reported for each species, the difference in use between plantations and other habitat types.

| Paper | Country | Response variable | Habitat use? | Species | Higher/lower use of plantations than native forest | Higher/lower use of plantations than native grassland | Higher/lower use of plantations than agricultural land |
| --- | --- | --- | --- | --- | --- | --- | --- |
| Acosta-Jamett and Simonetti, 2004 | Chile | Habitat selection | Yes | Leopardus guigna | Lower | NA | NA |
|  |  |  |  | Lycalopex culpaeus | Higher | NA | NA |
| Andrade-Núñez and Aide, 2010 | Uruguay | Presence/absence | Yes | Cerdocyon thous | NA | NA | NA |
|  |  |  |  | Conepatus chinga | No difference | Higher | NA |
|  |  |  |  | Leopardus geoffroyi | NA | NA | NA |
|  |  |  |  | Lycalopex gymnocercus | NA | NA | NA |
|  |  |  |  | Procyon cancrivorus | NA | NA | NA |
| Baker and Harris, 2006 | UK | Density | Yes | Vulpes vulpes | NA | NA | NA |
| Balakrishnan and Easa, 1986 | India | Abundance | Yes | Cuon alpinus | No difference | NA | NA |
|  |  |  |  | Melursus ursinus | Higher | NA | NA |
|  |  |  |  | Panthera pardus | No difference | NA | NA |
|  |  |  |  | Panthera tigris | No difference | NA | NA |
|  |  |  |  | Paradoxurus hermaphroditus | Lower | Higher | NA |
|  |  |  |  | Viverricula indica | No difference | NA | NA |
| Begotti et al., 2018 | Brazil | Presence/absence | Yes | Cerdocyon thous | NA | NA | NA |
|  |  |  |  | Chrysocyon brachyurus | NA | NA | NA |
|  |  |  |  | Eira barbara | NA | NA | NA |
|  |  |  |  | Galictis cuja | NA | NA | NA |
|  |  |  |  | Herpailurus yagouaroundi | NA | NA | NA |
|  |  |  |  | Leopardus pardalis | NA | NA | NA |
|  |  |  |  | Lontra longicaudis | NA | NA | NA |
|  |  |  |  | Nasua nasua | NA | NA | NA |
|  |  |  |  | Procyon cancrivorus | NA | NA | NA |
|  |  |  |  | Puma concolor | NA | NA | NA |
| Belden et al., 2007 | Malaysia | Presence | Yes | Herpestes brachyurus | NA | NA | NA |
|  |  |  |  | Herpestes semitorquatus | NA | NA | NA |
|  |  |  |  | Martes flavigula | NA | NA | NA |
|  |  |  |  | Mustela nudipes | NA | NA | NA |
|  |  |  |  | Paradoxurus hermaphroditus | NA | NA | NA |
|  |  |  |  | Viverra tangalunga | NA | NA | NA |
| Bojarska et al., 2021 | Poland | Rest site selection | Yes | Canis lupus | NA | NA | NA |
| Bonnington et al., 2009 | Tanzania | Presence/absence | Yes | Atilax paludinosus | NA | NA | NA |
|  |  |  |  | Canis mesomelas | NA | NA | NA |
|  |  |  |  | Civettictis civetta | NA | NA | NA |
|  |  |  |  | Panthera pardus | NA | NA | NA |
| Brandt and Lambin, 2007 | UK | Habitat selection | Yes | Mustela nivalis | NA | NA | NA |
| Calkoen et al., 2018 | Sweden | Land use intensity | Yes | Canis lupus | NA | NA | NA |
| Campos et al., 2018 | Brazil | Species diversity and distribution | Yes | Chrysocyon brachyurus | NA | NA | NA |
|  |  |  |  | Nasua nasua | NA | NA | NA |
|  |  |  |  | Puma concolor | NA | NA | NA |
| Caryl et al., 2012 | Scotland | Habitat selection | Yes | Martes martes | NA | Lower | Higher |
| Castro et al., 2022 | Portugal | Occurrence | Yes | Vulpes vulpes | Lower | NA | NA |
| Castro-Pastene et al., 2021 | Chile | Presence | Yes | Leopardus colocolo | NA | NA | NA |
| Chamberlain et al., 2002 | USA | Habitat selection | Yes | Procyon lotor | NA | NA | NA |
| Coelho et al., 2014 | Brazil | Presence/absence | Yes | Cerdocyon thous | Higher | Lower | NA |
| Cove et al., 2013 | Costa Rica | Occupancy | Yes | Eira barbara | NA | NA | NA |
|  |  |  |  | Leopardus pardalis | NA | NA | NA |
|  |  |  |  | Nasua narica | NA | NA | NA |
| Cravino and Brazeiro, 2021 | Uruguay | Capture rate | Yes | Procyon cancrivorus | NA | NA | NA |
| Cravino et al., 2023 | Uruguay | Species presence | Yes | Conepatus chinga | NA | NA | NA |
|  |  |  |  | Lycalopex gymnocercus | NA | NA | NA |
|  |  |  |  | Procyon cancrivorus | NA | NA | NA |
| Cresswell et al., 1989 | UK | Distribution | Yes | Meles meles | NA | NA | NA |
| Cruz et al., 2015 | Portugal | Occupancy | Yes | Martes foina | Lower | NA | NA |
|  |  |  |  | Meles meles | NA | NA | NA |
|  |  |  |  | Vulpes vulpes | Lower | NA | Lower |
| de Barros Ferraz et al., 2010 | Brazil | Distribution | Yes | Cerdocyon thous | No difference | NA | Lower |
| de Lima et al., 2013 | Brazil | Presence/absence | Yes | Panthera onca | NA | NA | NA |
| Dotta and Verdade, 2007 | Brazil | Presence/absence | Yes | Cerdocyon thous | NA | NA | NA |
|  |  |  |  | Chrysocyon brachyurus | NA | NA | NA |
|  |  |  |  | Conepatus chinga | NA | NA | NA |
|  |  |  |  | Eira barbara | NA | NA | NA |
|  |  |  |  | Galictis cuja | NA | NA | NA |
|  |  |  |  | Leopardus pardalis | NA | NA | NA |
|  |  |  |  | Nasua nasua | NA | NA | NA |
|  |  |  |  | Procyon cancrivorus | NA | NA | NA |
|  |  |  |  | Puma concolor | NA | NA | NA |
| Dotta and Verdade, 2009 | Brazil | Presence/absence | Yes | Herpailurus yagouaroundi | NA | NA | NA |
|  |  |  |  | Panthera onca | NA | NA | NA |
|  |  |  |  | Puma concolor | NA | NA | NA |
| Dotta and Verdade, 2011 | Brazil | Frequency of occurrence | Yes | Cerdocyon thous | NA | NA | NA |
|  |  |  |  | Chrysocyon brachyurus | NA | NA | NA |
|  |  |  |  | Conepatus chinga | NA | NA | NA |
|  |  |  |  | Eira barbara | NA | NA | NA |
|  |  |  |  | Galictis cuja | NA | NA | NA |
|  |  |  |  | Herpailurus yagouaroundi | NA | NA | NA |
|  |  |  |  | Leopardus pardalis | NA | NA | NA |
|  |  |  |  | Nasua nasua | NA | NA | NA |
|  |  |  |  | Procyon cancrivorus | NA | NA | NA |
|  |  |  |  | Puma concolor | NA | NA | NA |
| Eom et al., 2019 | South Korea | Habitat selection | Yes | Meles meles | NA | NA | NA |
|  |  |  |  | Nyctereutes procyonoides | NA | NA | NA |
| Escudero-Páez et al., 2018 | Chile | Frequency of occurrence | Yes | Conepatus chinga | No difference | NA | NA |
|  |  |  |  | Leopardus guigna | No difference | NA | NA |
|  |  |  |  | Lycalopex culpaeus | Higher | NA | NA |
|  |  |  |  | Puma concolor | NA | NA | NA |
| Fournier et al., 2007 | France | Habitat selection | Yes | Mustela lutreola | Lower | Lower | NA |
|  |  |  |  | Mustela putorius | Lower | No difference | No difference |
| Guerrero et al., 2006 | Chile | Habitat selection | Yes | Leopardus guigna | Lower | NA | NA |
|  |  |  |  | Lycalopex culpaeus | Higher | NA | NA |
|  |  |  |  | Lycalopex griseus | No difference | NA | NA |
| Homem et al., 2020 | Brazil | Presence | Yes | Cerdocyon thous | NA | NA | NA |
|  |  |  |  | Chrysocyon brachyurus | NA | NA | NA |
|  |  |  |  | Eira barbara | NA | NA | NA |
|  |  |  |  | Leopardus pardalis | NA | NA | NA |
|  |  |  |  | Nasua nasua | NA | NA | NA |
|  |  |  |  | Procyon cancrivorus | NA | NA | NA |
|  |  |  |  | Puma concolor | NA | NA | NA |
| Hwang et al., 2014 | South Korea | Abundance | Yes | Mustela sibirica | No difference | NA | NA |
|  |  |  |  | Nyctereutes procyonoides | Higher | NA | NA |
| Iezzi et al., 2018 | Argentina | Species presence | Yes | Cerdocyon thous | Higher | NA | NA |
|  |  |  |  | Eira barbara | No difference | NA | NA |
|  |  |  |  | Leopardus pardalis | Lower | NA | NA |
|  |  |  |  | Leopardus tigrinus | No difference | NA | NA |
|  |  |  |  | Nasua nasua | Lower | NA | NA |
|  |  |  |  | Panthera onca | No difference | NA | NA |
|  |  |  |  | Procyon cancrivorus | Lower | NA | NA |
|  |  |  |  | Puma concolor | No difference | NA | NA |
| Iezzi et al., 2020 | Argentina | Species presence | Yes | Chrysocyon brachyurus | NA | Lower | NA |
|  |  |  |  | Conepatus chinga | No difference | No difference | NA |
|  |  |  |  | Leopardus geoffroyi | Lower | No difference | NA |
|  |  |  |  | Leopardus pardalis | Lower | NA | NA |
|  |  |  |  | Lycalopex gymnocercus | No difference | Lower | NA |
|  |  |  |  | Nasua nasua | Lower | No difference | NA |
|  |  |  |  | Procyon cancrivorus | Lower | No difference | NA |
|  |  |  |  | Puma concolor | No difference | NA | NA |
| Iezzi et al., 2021 | Argentina | Assemblage | Yes | Cerdocyon thous | NA | NA | NA |
|  |  |  |  | Chrysocyon brachyurus | NA | NA | NA |
|  |  |  |  | Conepatus chinga | NA | NA | NA |
|  |  |  |  | Eira barbara | NA | NA | NA |
|  |  |  |  | Leopardus geoffroyi | NA | NA | NA |
|  |  |  |  | Leopardus guttulus | NA | NA | NA |
|  |  |  |  | Leopardus pardalis | NA | NA | NA |
|  |  |  |  | Leopardus wiedii | NA | NA | NA |
|  |  |  |  | Lycalopex gymnocercus | NA | NA | NA |
|  |  |  |  | Nasua nasua | NA | NA | NA |
|  |  |  |  | Panthera onca | NA | NA | NA |
|  |  |  |  | Procyon cancrivorus | NA | NA | NA |
|  |  |  |  | Puma concolor | NA | NA | NA |
| Jones and Pelton, 2003 | USA | Habitat selection | Yes | Ursus americanus | Lower | NA | NA |
| Jones et al., 2023 | Australia | Abundance | Yes | Dasyurus viverrinus | Higher | Lower | NA |
|  |  |  |  | Sarcophilus harrisii | Higher | NA | NA |
| Kaneko et al., 2006 | Japan | Habitat selection | Yes | Meles meles | NA | NA | NA |
| Kaneko et al., 2009 | Japan | Habitat selection, diet | Yes | Mustela itatsi | Higher | NA | NA |
| Karelus et al., 2018 | USA | Habitat selection | Yes | Ursus americanus | Lower | NA | NA |
| Katna et al., 2022 | India | Habitat selection | Yes | Canis aureus | NA | Lower | Lower |
|  |  |  |  | Felis chaus | NA | Lower | Lower |
|  |  |  |  | Vulpes bengalensis | NA | Higher | Higher |
| Kheswa et al., 2018 | South Africa | Occupancy | Yes | Mellivora capensis | Higher | NA | NA |
| Lantschner et al., 2012 | Argentina | Habitat selection | Yes | Conepatus chinga | Higher | NA | NA |
|  |  |  |  | Lycalopex culpaeus | Lower | NA | NA |
|  |  |  |  | Puma concolor | No difference | NA | NA |
| Linnell et al., 2017 | USA | Habitat selection | Yes | Mustela erminea | NA | NA | NA |
| Llaneza et al., 2016 | Spain | Rest site selection | Yes | Canis lupus | Higher | Higher | Higher |
| Lyall, 2017 | Australia | Abundance | Yes | Dasyurus maculatus | Lower | NA | NA |
|  |  |  |  | Sarcophilus harrisii | Lower | NA | NA |
| Lyra-Jorge et al., 2008 | Brazil | Presence/absence | Yes | Chrysocyon brachyurus | Higher | NA | NA |
|  |  |  |  | Conepatus semistriatus | Lower | NA | NA |
|  |  |  |  | Eira barbara | Higher | NA | NA |
|  |  |  |  | Leopardus pardalis | Lower | NA | NA |
|  |  |  |  | Puma concolor | NA | NA | NA |
| Lyra-Jorge et al., 2010 | Brazil | Presence/absence | Yes | Chrysocyon brachyurus | NA | NA | NA |
|  |  |  |  | Eira barbara | NA | NA | NA |
|  |  |  |  | Herpailurus yagouaroundi | NA | NA | NA |
|  |  |  |  | Leopardus pardalis | Lower | NA | NA |
|  |  |  |  | Puma concolor | NA | NA | NA |
| Manzo et al., 2018 | Italy | Home range | Yes | Martes martes | NA | NA | Lower |
| Marques and Fabián, 2013 | Brazil | Presence | Yes | Chrysocyon brachyurus | NA | NA | NA |
| Mazzolli, 2010 | Brazil | Density | Yes | Puma concolor | No difference | NA | NA |
| McShea et al., 2009 | Malaysia | Occupancy | Yes | Helarctos malayanus | NA | NA | NA |
|  |  |  |  | Hemigalus derbyanus | NA | NA | NA |
|  |  |  |  | Herpestes brachyurus | NA | NA | NA |
|  |  |  |  | Herpestes semitorquatus | NA | NA | NA |
|  |  |  |  | Martes flavigula | NA | NA | NA |
|  |  |  |  | Paradoxurus hermaphroditus | NA | NA | NA |
|  |  |  |  | Pardofelis marmorata | NA | NA | NA |
|  |  |  |  | Prionailurus bengalensis | NA | NA | NA |
|  |  |  |  | Viverra tangalunga | NA | NA | NA |
| Michalski et al., 2006 | Brazil | Habitat use | Yes | Cerdocyon thous | No difference | No difference | NA |
|  |  |  |  | Eira barbara | No difference | No difference | NA |
|  |  |  |  | Herpailurus yagouaroundi | NA | NA | NA |
| Moreira-Arce et al., 2015 | Chile | Diet | No | Leopardus guigna | NA | NA | NA |
|  |  |  |  | Lycalopex culpaeus | NA | NA | NA |
|  |  |  |  | Lycalopex fulvipes | NA | NA | NA |
|  |  |  |  | Lycalopex griseus | NA | NA | NA |
| Moreira-Arce et al., 2016 | Chile | Occupancy | Yes | Leopardus guigna | Lower | NA | NA |
|  |  |  |  | Lycalopex culpaeus | Higher | NA | NA |
|  |  |  |  | Lycalopex fulvipes | No difference | NA | NA |
|  |  |  |  | Lycalopex griseus | No difference | NA | NA |
| Munoz and Murua, 1990 | Chile | Presence/absence | Yes | Lycalopex culpaeus | NA | NA | NA |
| Ng et al., 2021 | Malaysia | Presence/absence | Yes | Helarctos malayanus | NA | NA | NA |
|  |  |  |  | Martes flavigula | NA | NA | NA |
|  |  |  |  | Mydaus javanensis | NA | NA | NA |
|  |  |  |  | Prionailurus bengalensis | NA | NA | NA |
| O’Mahony et al., 1999 | UK | Diet | No | Vulpes vulpes | NA | NA | NA |
| O’Mahony, 2014 | Ireland | Home range | Yes | Martes martes | NA | NA | NA |
| Palomares et al., 2000 | Spain | Habitat selection | Yes | Lynx pardinus | NA | Lower | Higher |
| Paolino et al., 2018 | Brazil | Abundance | Yes | Leopardus pardalis | Lower | NA | NA |
| Parry et al., 2009 | Brazil | Hunting success | No | Leopardus pardalis | NA | NA | NA |
|  |  |  |  | Nasua nasua | NA | NA | NA |
|  |  |  |  | Panthera onca | NA | NA | NA |
| Paviolo et al., 2018 | Argentina | Occupancy | Yes | Panthera onca | No difference | NA | NA |
|  |  |  |  | Puma concolor | No difference | NA | NA |
| Pedersen et al., 2009 | Norway | Predation rates | No | Martes martes | NA | NA | NA |
|  |  |  |  | Mustela erminea | NA | NA | NA |
|  |  |  |  | Vulpes vulpes | NA | NA | NA |
| Pedersen et al., 2010 | Norway | Species distribution | Yes | Lynx lynx | NA | NA | NA |
|  |  |  |  | Martes martes | Lower | NA | NA |
|  |  |  |  | Mustela erminea | NA | NA | NA |
|  |  |  |  | Mustela nivalis | NA | NA | NA |
|  |  |  |  | Vulpes vulpes | Lower | NA | NA |
| Pereira et al., 2012 | Portugal | Habitat selection | Yes | Genetta genetta | Higher | NA | Higher |
|  |  |  |  | Martes foina | Lower | NA | NA |
|  |  |  |  | Vulpes vulpes | NA | NA | NA |
| Piña et al., 2019 | Brazil | Species presence | Yes | Cerdocyon thous | NA | NA | NA |
| Pita et al., 2009 | Portugal | Abundance | Yes | Herpestes ichneumon | NA | NA | No difference |
|  |  |  |  | Lutra lutra | NA | NA | No difference |
|  |  |  |  | Meles meles | NA | NA | Higher |
|  |  |  |  | Mustela nivalis | NA | NA | No difference |
|  |  |  |  | Vulpes vulpes | NA | NA | Higher |
| Pita et al., 2020 | Portugal | Occupancy | Yes | Meles meles | Higher | NA | Higher |
| Ramesh and Downs, 2015 | South Africa | Occupancy | Yes | Herpestes ichneumon | Higher | NA | No difference |
| Ramesh et al., 2016a | South Africa | Home range | Yes | Leptailurus serval | Lower | Lower | Higher |
| Ramesh et al., 2016b | South Africa | Occupancy | Yes | Aonyx capensis | NA | NA | NA |
|  |  |  |  | Atilax paludinosus | No difference | No difference | NA |
|  |  |  |  | Canis adustus | Higher | Higher | NA |
|  |  |  |  | Crocuta crocuta | No difference | No difference | NA |
|  |  |  |  | Galerella sanguineus | Lower | NA | NA |
|  |  |  |  | Genetta tigrina | No difference | No difference | NA |
|  |  |  |  | Ichneumia albicauda | Lower | NA | NA |
|  |  |  |  | Ictonyx striatus | Lower | NA | NA |
|  |  |  |  | Leptailurus serval | NA | Lower | NA |
|  |  |  |  | Lycaon pictus | Lower | NA | NA |
|  |  |  |  | Mellivora capensis | No difference | Higher | NA |
|  |  |  |  | Mungos mungo | NA | NA | NA |
|  |  |  |  | Panthera pardus | NA | NA | NA |
|  |  |  |  | Rhynchogale melleri | Lower | NA | NA |
| Ramesh et al., 2017a | South Africa | Habitat selection | Yes | Caracal caracal | Higher | Higher | Higher |
| Ramesh et al., 2017b | South Africa | Density | Yes | Panthera pardus | Lower | NA | NA |
| Revilla et al., 2000 | Spain | Habitat selection | Yes | Meles meles | NA | Lower | NA |
| Rhim et al., 2015 | South Korea | Abundance | Yes | Meles meles | No difference | NA | NA |
|  |  |  |  | Mustela sibirica | No difference | NA | NA |
|  |  |  |  | Nyctereutes procyonoides | No difference | NA | NA |
| Rosalino et al., 2004 | Portugal | Density | Yes | Meles meles | Lower | NA | NA |
| Santos and Beier, 2008 | Portugal | Habitat selection | Yes | Meles meles | Lower | NA | NA |
| Sato et al., 2008 | Japan | Home range and habitat selection | Yes | Ursus arctos | Lower | NA | NA |
| Simonetti et al., 2013 | Chile | Frequency of occurrence | Yes | Conepatus chinga | NA | NA | NA |
|  |  |  |  | Leopardus guigna | NA | NA | NA |
|  |  |  |  | Lycalopex culpaeus | NA | NA | NA |
| Sompud et al., 2016 | Malaysia | Density | Yes | Arctogalidia trivirgata | NA | NA | NA |
|  |  |  |  | Paradoxurus hermaphroditus | NA | NA | NA |
| Son et al., 2017 | South Korea | Abundance | Yes | Meles meles | NA | NA | NA |
|  |  |  |  | Mustela sibirica | NA | NA | NA |
|  |  |  |  | Nyctereutes procyonoides | NA | NA | NA |
| Stratman et al., 2001 | USA | Habitat selection | Yes | Ursus americanus | NA | NA | Higher |
| Sunarto et al., 2012 | Indonesia | Occupancy | Yes | Panthera tigris | Lower | NA | NA |
| Takahata et al., 2013 | Japan | Habitat selection | Yes | Ursus thibetanus | Lower | NA | NA |
| Tanigawa et al., 2022 | Japan | Occupancy | Yes | Martes melampus | No difference | NA | NA |
|  |  |  |  | Meles anakuma | Lower | NA | NA |
|  |  |  |  | Nyctereutes procyonoides | Lower | NA | NA |
|  |  |  |  | Paguma larvata | No difference | NA | NA |
|  |  |  |  | Vulpes vulpes | Lower | NA | NA |
| Timo et al., 2014 | Brazil | Presence/absence | Yes | Canis lupus | NA | NA | NA |
|  |  |  |  | Cerdocyon thous | NA | NA | NA |
|  |  |  |  | Chrysocyon brachyurus | NA | NA | NA |
|  |  |  |  | Eira barbara | NA | NA | NA |
|  |  |  |  | Leopardus tigrinus | NA | NA | NA |
|  |  |  |  | Nasua nasua | NA | NA | NA |
|  |  |  |  | Puma concolor | NA | NA | NA |
| Tófoli et al., 2009 | Brazil | Diet | No | Herpailurus yagouaroundi | NA | NA | NA |
| Tomita and Hiura, 2021 | Japan | Foraging habitat selection | Yes | Ursus arctos | Higher | NA | NA |
| Tsujino and Yumoto, 2014 | Japan | Habitat selection | Yes | Martes melampus | Higher | NA | NA |
|  |  |  |  | Paguma larvata | Higher | NA | NA |
|  |  |  |  | Ursus thibetanus | Higher | NA | NA |
| Twining et al., 2019 | Ireland | Diet | No | Martes martes | NA | NA | NA |
| Twining et al., 2020 | Ireland | Habitat selection | Yes | Martes martes | Lower | No difference | Higher |
| Twining et al., 2022 | Ireland | Occupancy | Yes | Martes martes | Higher | NA | NA |
| Vanak and Gompper, 2010 | India | Habitat selection | Yes | Vulpes bengalensis | NA | Lower | Higher |
| Vergara et al., 2016 | Spain | Presence | Yes | Martes foina | NA | NA | Lower |
|  |  |  |  | Martes martes | Lower | NA | NA |
| Virgós et al., 2002 | Spain | Habitat selection | Yes | Felis silvestris | Lower | NA | NA |
|  |  |  |  | Genetta genetta | Higher | NA | NA |
|  |  |  |  | Martes foina | Lower | NA | NA |
|  |  |  |  | Meles meles | Lower | NA | NA |
|  |  |  |  | Vulpes vulpes | Lower | NA | NA |
| Willcox et al., 2012 | Vietnam | Presence | Yes | Arctogalidia trivirgata | NA | NA | NA |
| Wong et al., 2022 | Malaysia | Occupancy | Yes | Aonyx cinereus | No difference | NA | NA |
|  |  |  |  | Catopuma badia | No difference | NA | NA |
|  |  |  |  | Diplogale hosei | Lower | NA | NA |
|  |  |  |  | Helarctos malayanus | No difference | NA | NA |
|  |  |  |  | Hemigalus derbyanus | No difference | NA | NA |
|  |  |  |  | Martes flavigula | No difference | NA | NA |
|  |  |  |  | Paguma larvata | No difference | NA | NA |
|  |  |  |  | Paradoxurus hermaphroditus | Higher | NA | NA |
|  |  |  |  | Pardofelis marmorata | No difference | NA | NA |
|  |  |  |  | Prionailurus bengalensis | No difference | NA | NA |
|  |  |  |  | Viverra tangalunga | No difference | NA | NA |
| Yaap et al., 2016 | Indonesia | Habitat selection | Yes | Helarctos malayanus | NA | NA | NA |
|  |  |  |  | Herpestes brachyurus | NA | NA | NA |
|  |  |  |  | Panthera tigris | NA | NA | NA |
|  |  |  |  | Paradoxurus hermaphroditus | NA | NA | NA |
|  |  |  |  | Viverra tangalunga | NA | NA | NA |
| Yamada and Fujioka, 2010 | Japan | Presence/absence | Yes | Ursus thibetanus | NA | NA | NA |
| Zabala et al., 2005 | Spain | Habitat selection | Yes | Mustela putorius | Lower | Lower | NA |
| Zhang et al., 2019 | China | Habitat selection | Yes | Martes flavigula | Lower | NA | NA |
|  |  |  |  | Mustela sibirica | Lower | NA | NA |
|  |  |  |  | Paguma larvata | Lower | NA | NA |
|  |  |  |  | Prionailurus bengalensis | NA | NA | NA |
|  |  |  |  | Ursus thibetanus | Lower | NA | NA |
| Zuberogoitia et al., 2002 | Spain | Habitat selection | Yes | Genetta genetta | Lower | NA | NA |
| Zúñiga et al., 2009 | Spain | Habitat selection | Yes | Galictis cuja | Higher | Higher | NA |
|  |  |  |  | Leopardus guigna | No difference | NA | NA |
|  |  |  |  | Lycalopex griseus | Higher | Higher | NA |
|  |  |  |  | Puma concolor | Higher | Higher | NA |

**Appendix S2:** Summary of papers reporting carnivores using harvested native forest, including the country in which the research was conducted, the harvesting method, the response variable considered, whether the response variable fit our definition of ‘habitat use’, the species recorded, and where reported for each species, the difference in use between harvested and unharvested forest.

| Paper | Country | Harvesting type | Response variable | Habitat use? | Species | Higher use of harvested/  unharvested forest |
| --- | --- | --- | --- | --- | --- | --- |
| Andruskiw et al., 2008 | Canada | Mixed | Hunting success | No | Martes americana | Harvested forest |
| Augeri, 2005 | Indonesia | Mixed | Density | Yes | Helarctos malayanus | Unharvested forest |
| Ausilio et al., 2022 | Norway and Sweden | Clearfelling | Hunting success | No | Canis lupus | Harvested forest |
| Bahaa-el-din et al., 2016 | Gabon | Unspecified | Density | Yes | Caracal aurata | No difference |
| Belcher, 2003 | Australia | Partial harvesting | Population dynamics | No | Dasyurus maculatus | No difference |
| Belcher, 2008 | Australia | Partial harvesting | Habitat use | Yes | Dasyurus maculatus | Unharvested forest |
| Bernard et al., 2013 | Malaysia | Unspecified | Species presence | Yes | Herpestes semitorquatus | NA |
|  |  |  |  |  | Lutrogale perspicillata | NA |
|  |  |  |  |  | Neofelis diardi | NA |
|  |  |  |  |  | Paguma larvata | NA |
|  |  |  |  |  | Pardofelis marmorata | NA |
|  |  |  |  |  | Prionailurus bengalensis | NA |
|  |  |  |  |  | Viverra tangalunga | NA |
| Bobo et al., 2017 | Cameroon | Unspecified | Abundance | Yes | Civettictis civetta | NA |
|  |  |  |  |  | Herpestes naso | NA |
| Boisjoly et al., 2010 | Canada | Clearfelling | Habitat use | Yes | Canis latrans | Harvested forest |
| Bojarska et al., 2017 | Poland | Unspecified | Kill site locations | No | Canis lupus | NA |
| Bourbonnais, 2013 | Canada | Clearfelling | Health | No | Ursus arctos | NA |
| Bowman et al., 2010 | Canada | Unspecified | Occurrence | Yes | Canis lupus | Harvested forest |
|  |  |  |  |  | Gulo gulo | Unharvested forest |
| Brodie and Giordano, 2011 | Malaysia | Partial harvesting | Presence | Yes | Arctictis binturong | NA |
|  |  |  |  |  | Hemigalus derbyanus | NA |
|  |  |  |  |  | Martes flavigula | NA |
|  |  |  |  |  | Paradoxurus hermaphroditus | NA |
| Brodie et al., 2015 | Malaysia | Partial harvesting | Abundance | Yes | Arctictis binturong | NA |
|  |  |  |  |  | Helarctos malayanus | No difference |
|  |  |  |  |  | Hemigalus derbyanus | No difference |
|  |  |  |  |  | Neofelis diardi | Unharvested forest |
|  |  |  |  |  | Paradoxurus hermaphroditus | NA |
| Broquet et al., 2006 | Canada | Unspecified | Gene flow | No | Martes americana | NA |
| Bull and Heater, 2000 | USA | Unspecified | Rest site selection | Yes | Martes americana | NA |
| Bull et al., 2005 | USA | Partial harvesting | Habitat use | Yes | Martes americana | Harvested forest |
| Buskirk et al., 1996 | China | Partial harvesting | Habitat use | Yes | Martes zibellina | NA |
| Carvalho Jr et al., 2021 | Brazil | Reduced impact logging | Occupancy | Yes | Atelocynus microtis | No difference |
|  |  |  |  |  | Eira barbara | No difference |
|  |  |  |  |  | Herpailurus yagouaroundi | No difference |
|  |  |  |  |  | Leopardus pardalis | No difference |
|  |  |  |  |  | Leopardus wiedii | No difference |
|  |  |  |  |  | Nasua nasua | No difference |
|  |  |  |  |  | Panthera onca | No difference |
|  |  |  |  |  | Procyon cancrivorus | No difference |
|  |  |  |  |  | Puma concolor | No difference |
| Chanchani et al., 2016 | India | Unspecified | Occupancy | Yes | Panthera tigris | No difference |
| Chapin et al., 1998 | USA | Unspecified | Habitat use | Yes | Martes americana | Unharvested forest |
| Clements et al., 2021 | Malaysia | Unspecified | Detection frequency | Yes | Arctictis binturong | NA |
|  |  |  |  |  | Catopuma temminckii | NA |
|  |  |  |  |  | Cuon alpinus | NA |
|  |  |  |  |  | Helarctos malayanus | NA |
|  |  |  |  |  | Hemigalus derbyanus | NA |
|  |  |  |  |  | Herpestes urva | NA |
|  |  |  |  |  | Martes flavigula | NA |
|  |  |  |  |  | Mustela nudipes | NA |
|  |  |  |  |  | Neofelis nebulosa | NA |
|  |  |  |  |  | Paguma larvata | NA |
|  |  |  |  |  | Panthera pardus | NA |
|  |  |  |  |  | Panthera tigris | NA |
|  |  |  |  |  | Paradoxurus hermaphroditus | NA |
|  |  |  |  |  | Pardofelis marmorata | NA |
|  |  |  |  |  | Prionailurus bengalensis | NA |
|  |  |  |  |  | Prionodon linsang | NA |
|  |  |  |  |  | Viverra tangalunga | NA |
| Colón, 2002 | Malaysia | Partial harvesting | Density | Yes | Viverra tangalunga | Unharvested forest |
| Constantinou, 2021 | Canada | Mixed | Habitat use | Yes | Canis lupus | NA |
|  |  |  |  |  | Gulo gulo | NA |
|  |  |  |  |  | Lynx canadensis | NA |
|  |  |  |  |  | Lynx rufus | NA |
|  |  |  |  |  | Martes americana | NA |
|  |  |  |  |  | Mephitis mephitis | NA |
|  |  |  |  |  | Mustela erminea | NA |
|  |  |  |  |  | Mustela nivalis | NA |
|  |  |  |  |  | Puma concolor | NA |
|  |  |  |  |  | Ursus arctos | NA |
|  |  |  |  |  | Vulpes vulpes | NA |
| Crimmins et al., 2012 | USA | Clearfelling | Habitat use | Yes | Canis latrans | Harvested forest |
| Di Bitetti et al., 2008 | Argentina | Unspecified | Density | Yes | Leopardus pardalis | Unharvested forest |
| Dijak and Thompson, 2000 | USA | Clearfelling | Abundance | Yes | Mephitis mephitis | No difference |
|  |  |  |  |  | Procyon lotor | No difference |
| Evans and Mortelliti, 2022 | USA | Unspecified | Occupancy | Yes | Martes americana | Unharvested forest |
|  |  |  |  |  | Martes pennanti | Unharvested forest |
| Evans and Mortelliti, 2022 | USA | Partial harvesting | Occupancy | Yes | Mustela erminea | Harvested forest |
|  |  |  |  |  | Mustela frenata | Harvested forest |
| Fisher et al., 2013 | Canada | Unspecified | Abundance/occupancy | Yes | Gulo gulo | Unharvested forest |
| Flynn et al., 2011 | Australia | Clearfelling | Abundance | Yes | Dasyurus maculatus | No difference |
|  |  |  |  |  | Dasyurus viverrinus | Harvested forest |
| Fuller and Harrison, 2005 | USA | Mixed | Habitat selection | Yes | Martes americana | Harvested forest |
| Fuller and Harrison, 2010 | USA | Mixed | Habitat use | Yes | Lynx canadensis | Harvested forest |
| Gagné et al., 2016 | Canada | Clearfelling | Occurrence | Yes | Canis lupus | Harvested forest |
| Gerber et al., 2012 | Madagascar | Partial harvesting | Occupancy | Yes | Cryptoprocta ferox | No difference |
|  |  |  |  |  | Eupleres goudotii | NA |
|  |  |  |  |  | Fossa fossana | Unharvested forest |
|  |  |  |  |  | Galidia elegans | NA |
|  |  |  |  |  | Galidictis fasciata | NA |
| Gervasi et al., 2013 | Scandinavia | Clearfelling | Predation events | No | Canis lupus | NA |
| Goszczyński et al., 2007 | Poland | Unspecified | Habitat use | Yes | Martes foina | NA |
|  |  |  |  |  | Martes martes | Unharvested forest |
| Granados et al., 2016 | Malaysia | Partial harvesting | Abundance | Yes | Hemigalus derbyanus | No difference |
|  |  |  |  |  | Viverra tangalunga | No difference |
| Guharajan et al., 2021 | Malaysia | Mixed | Occupancy | Yes | Helarctos malayanus | NA |
| Gulsby et al., 2017 | USA | Clearfelling | Predation events | No | Canis latrans | NA |
| Gurarie et al., 2011 | Finland | Clearfelling | Habitat use | Yes | Canis lupus | Harvested forest |
| Gurtler, 2020 | USA | Partial harvesting | Community composition | Yes | Canis lupus | NA |
|  |  |  |  |  | Martes americana | NA |
|  |  |  |  |  | Procyon lotor | NA |
|  |  |  |  |  | Ursus americanus | NA |
| Gutiérrez-Granados and Dirzo, 2021 | Mexico | Partial harvesting | Abundance | Yes | Conepatus semistriatus | No difference |
|  |  |  |  |  | Herpailurus yagouaroundi | No difference |
|  |  |  |  |  | Nasua narica | Unharvested forest |
|  |  |  |  |  | Panthera onca | NA |
|  |  |  |  |  | Urocyon cinereoargenteus | Unharvested forest |
| Hansson, 1994 | Sweden | Clearfelling | Abundance | Yes | Martes martes | Unharvested forest |
|  |  |  |  |  | Mustela erminea | Harvested forest |
|  |  |  |  |  | Mustela nivalis | Harvested forest |
|  |  |  |  |  | Vulpes vulpes | No difference |
| Happe et al., 2020 | USA | Unspecified | Occupancy | Yes | Martes pennanti | Unharvested forest |
| Hargis et al., 1999 | USA | Clearfelling | Capture rate | Yes | Martes americana | NA |
| Haysom et al., 2021 | Malaysia | Unspecified | Community composition | Yes | Arctictis binturong | NA |
|  |  |  |  |  | Arctogalidia trivirgata | NA |
|  |  |  |  |  | Helarctos malayanus | NA |
|  |  |  |  |  | Hemigalus derbyanus | NA |
|  |  |  |  |  | Herpestes brachyurus | NA |
|  |  |  |  |  | Herpestes semitorquatus | NA |
|  |  |  |  |  | Martes flavigula | NA |
|  |  |  |  |  | Mydaus javanensis | NA |
|  |  |  |  |  | Neofelis diardi | NA |
|  |  |  |  |  | Paguma larvata | NA |
|  |  |  |  |  | Paradoxurus hermaphroditus | NA |
|  |  |  |  |  | Pardofelis marmorata | NA |
|  |  |  |  |  | Prionailurus bengalensis | NA |
|  |  |  |  |  | Viverra tangalunga | NA |
| Hearn et al., 2010 | Canada | Clearfelling | Habitat use | Yes | Martes americana | NA |
| Hearn et al., 2016 | Malaysia | Partial harvesting | Density and distribution | Yes | Pardofelis marmorata | No difference |
| Hearn et al., 2019 | Malaysia | Partial harvesting | Abundance | Yes | Neofelis diardi | NA |
| Heydon and Bulloh, 1996 | Malaysia | Partial harvesting | Abundance | Yes | Arctictis binturong | NA |
|  |  |  |  |  | Hemigalus derbyanus | Unharvested forest |
|  |  |  |  |  | Paradoxurus hermaphroditus | NA |
|  |  |  |  |  | Viverra tangalunga | Unharvested forest |
| Holbrook et al., 2017 | USA | Unspecified | Habitat use | Yes | Lynx canadensis | Unharvested forest |
| Holbrook et al., 2018 | USA | Mixed | Habitat use | Yes | Lynx canadensis | NA |
| Holbrook et al., 2019 | USA | Unspecified | Reproductive success | No | Lynx canadensis | Unharvested forest |
| Homkes, 2021 | USA | Unspecified | Kill sites | No | Canis lupus | NA |
| Houle et al., 2010 | Canada | Clearfelling | Habitat selection | Yes | Canis lupus | NA |
| Hoving et al., 2004 | USA | Mixed | Habitat use | Yes | Lynx canadensis | NA |
| Imron et al., 2011 | Indonesia | Partial harvesting | Probability of extinction | No | Panthera tigris | Unharvested forest |
| Jamhuri et al., 2018 | Malaysia | Partial harvesting | Species occurrence | Yes | Helarctos malayanus | NA |
| Jati et al., 2018 | Indonesia | Reduced impact logging | Species presence | Yes | Arctictis binturong | NA |
|  |  |  |  |  | Catopuma badia | NA |
|  |  |  |  |  | Cynogale bennettii | NA |
|  |  |  |  |  | Diplogale hosei | NA |
|  |  |  |  |  | Helarctos malayanus | NA |
|  |  |  |  |  | Hemigalus derbyanus | NA |
|  |  |  |  |  | Herpestes brachyurus | Harvested forest |
|  |  |  |  |  | Herpestes semitorquatus | NA |
|  |  |  |  |  | Martes flavigula | Harvested forest |
|  |  |  |  |  | Mustela nudipes | NA |
|  |  |  |  |  | Neofelis diardi | NA |
|  |  |  |  |  | Paguma larvata | Harvested forest |
|  |  |  |  |  | Paradoxurus hermaphroditus | No difference |
|  |  |  |  |  | Pardofelis marmorata | NA |
|  |  |  |  |  | Prionailurus bengalensis | NA |
|  |  |  |  |  | Prionailurus planiceps | NA |
|  |  |  |  |  | Prionodon linsang | NA |
|  |  |  |  |  | Viverra tangalunga | Unharvested forest |
| Jennings et al., 2006 | Indonesia | Partial harvesting | Home range and activity | Yes | Viverra tangalunga | Unharvested forest |
| Jiang et al., 2017 | China | Unspecified | Abundance | Yes | Panthera pardus | Unharvested forest |
|  |  |  |  |  | Panthera tigris | Unharvested forest |
| Jokinen et al., 2019 | Canada | Unspecified | Den sites | Yes | Gulo gulo | NA |
| Jones and Pelton, 2003 | USA | Clearfelling | Habitat use | Yes | Ursus americanus | Harvested forest |
| Jones et al., 2023 | Australia | Partial harvesting | Abundance | Yes | Dasyurus viverrinus | No difference |
|  |  |  |  |  | Sarcophilus harrisii | No difference |
| Kaartinen et al., 2009 | Finland | Clearfelling | Depredation risk to sheep | No | Canis lupus | NA |
| Kaartinen et al., 2015 | Finland | Unspecified | Habitat use | Yes | Canis lupus | NA |
| Kauhala and Ihalainen, 2014 | Finland | Clearfelling | Diet | No | Meles meles | NA |
| Kays et al., 2008 | USA | Partial harvesting | Abundance | Yes | Canis latrans | NA |
| Kertson and Marzluff, 2011 | USA | Clearfelling | Habitat use | Yes | Puma concolor | Harvested forest |
| Kitamura et al., 2010 | Thailand | Unspecified | Species presence | Yes | Arctictis binturong | NA |
|  |  |  |  |  | Catopuma temminckii | NA |
|  |  |  |  |  | Hemigalus derbyanus | NA |
|  |  |  |  |  | Martes flavigula | NA |
|  |  |  |  |  | Paguma larvata | NA |
|  |  |  |  |  | Panthera pardus | NA |
|  |  |  |  |  | Panthera tigris | NA |
|  |  |  |  |  | Paradoxurus hermaphroditus | NA |
|  |  |  |  |  | Prionailurus bengalensis | NA |
|  |  |  |  |  | Prionodon linsang | NA |
| Kordosky et al., 2021 | USA | Thinning | Home range | Yes | Martes pennanti | NA |
| Kortello et al., 2019 | Canada | Unspecified | Occupancy | Yes | Gulo gulo | Unharvested forest |
| Kosterman et al., 2018 | USA | Unspecified | Reproductive success | No | Lynx canadensis | Unharvested forest |
| Krebs et al., 2007 | Canada | Unspecified | Habitat use | Yes | Gulo gulo | NA |
| Krofel et al., 2017 | Slovenia | Unspecified | Scent marking | No | Lynx lynx | NA |
| Kunkel and Pletscher, 2000 | Canada | Clearfelling | Kill sites | No | Canis lupus | NA |
| Kuzyk et al., 2004 | Canada | Clearfelling | Habitat use | Yes | Canis lupus | No difference |
| Laidlaw, 2000 | Malaysia | Partial harvesting | Species presence | Yes | Helarctos malayanus | NA |
|  |  |  |  |  | Martes flavigula | NA |
|  |  |  |  |  | Neofelis nebulosa | NA |
|  |  |  |  |  | Panthera tigris | NA |
| Laufer et al., 2015 | Brazil | Reduced impact logging | Species presence | Yes | Eira barbara | NA |
|  |  |  |  |  | Panthera onca | NA |
|  |  |  |  |  | Puma concolor | NA |
| Lavoie et al., 2019 | Canada | Thinning | Abundance | Yes | Martes americana | Unharvested forest |
| Lesmeister et al., 2013 | USA | Unspecified | Occupancy | Yes | Spilogale putorius | NA |
| Lesmerises et al., 2012 | Canada | Clearfelling | Habitat use | Yes | Canis lupus | NA |
| Lhoest, 2020 | Cameroon | Partial harvesting | Species presence | Yes | Atilax paludinosus | NA |
|  |  |  |  |  | Bdeogale nigripes | NA |
|  |  |  |  |  | Crossarchus platycephalus | NA |
|  |  |  |  |  | Genetta servalina | NA |
|  |  |  |  |  | Nandinia binotata | NA |
| Linkie et al., 2003 | Indonesia | Unspecified | Distribution | Yes | Panthera tigris | NA |
| Linkie et al., 2007 | Indonesia | Partial harvesting | Occupancy | Yes | Helarctos malayanus | Harvested forest |
| Linkie et al., 2008 | Indonesia | Partial harvesting | Density | Yes | Panthera tigris | Unharvested forest |
| Lisgo, 1999 | Canada | Clearfelling | Habitat use | Yes | Mustela erminea | NA |
| Lomolino and Perault, 2000 | USA | Clearfelling | Species presence | Yes | Lynx rufus | NA |
|  |  |  |  |  | Mustela erminea | NA |
|  |  |  |  |  | Mustela frenata | NA |
|  |  |  |  |  | Spilogale putorius | NA |
|  |  |  |  |  | Ursus americanus | NA |
| Lomolino and Perault, 2001 | USA | Clearfelling | Distribution | Yes | Mustela erminea | Harvested forest |
|  |  |  |  |  | Mustela frenata | NA |
|  |  |  |  |  | Spilogale putorius | Harvested forest |
|  |  |  |  |  | Ursus americanus | Unharvested forest |
| Lopes and Ferrari, 2000 | Brazil | Unspecified | Species presence | Yes | Eira barbara | NA |
|  |  |  |  |  | Nasua nasua | NA |
|  |  |  |  |  | Speothos venaticus | NA |
| Luskin, 2016 | Indonesia | Partial harvesting | Density | Yes | Panthera tigris | Unharvested forest |
| Magintan et al., 2017 | Malaysia | Unspecified | Species presence | Yes | Aonyx cinereus | NA |
|  |  |  |  |  | Catopuma temminckii | NA |
|  |  |  |  |  | Helarctos malayanus | NA |
|  |  |  |  |  | Lutrogale perspicillata | NA |
|  |  |  |  |  | Panthera tigris | NA |
| Maiwald et al., 2021 | Malaysia | Reduced impact logging | Occupancy | Yes | Catopuma badia | No difference |
|  |  |  |  |  | Neofelis diardi | No difference |
|  |  |  |  |  | Pardofelis marmorata | No difference |
|  |  |  |  |  | Prionailurus bengalensis | No difference |
| Malcolm et al., 2004 | Canada | Clearfelling | Habitat suitability | Yes | Lynx canadensis | Unharvested forest |
|  |  |  |  |  | Martes americana | No difference |
|  |  |  |  |  | Procyon lotor | Unharvested forest |
|  |  |  |  |  | Ursus americanus | Unharvested forest |
| Mangas and Rodríguez-Estival, 2010 | Spain | Mixed | Abundance | Yes | Vulpes vulpes | NA |
| Mathai et al., 2010 | Malaysia | Reduced impact logging | Presence | Yes | Arctictis binturong | NA |
|  |  |  |  |  | Helarctos malayanus | NA |
|  |  |  |  |  | Herpestes brachyurus | NA |
|  |  |  |  |  | Martes flavigula | NA |
|  |  |  |  |  | Mustela nudipes | NA |
|  |  |  |  |  | Viverra tangalunga | NA |
| Mathai et al., 2014 | Malaysia | Unspecified | Presence | Yes | Catopuma badia | NA |
|  |  |  |  |  | Neofelis diardi | NA |
|  |  |  |  |  | Pardofelis marmorata | NA |
|  |  |  |  |  | Prionailurus bengalensis | NA |
| Mathai et al., 2017 | Malaysia | Mixed | Occupancy | Yes | Diplogale hosei | Unharvested forest |
|  |  |  |  |  | Hemigalus derbyanus | Unharvested forest |
|  |  |  |  |  | Herpestes brachyurus | Harvested forest |
|  |  |  |  |  | Martes flavigula | Unharvested forest |
|  |  |  |  |  | Paguma larvata | NA |
|  |  |  |  |  | Prionailurus bengalensis | NA |
|  |  |  |  |  | Viverra tangalunga | Harvested forest |
| Mathur et al., 2011 | India | Unspecified | Abundance | Yes | Panthera tigris | NA |
| Matthews and Matthews, 2006 | Cameroon | Unspecified | Presence/absence | Yes | Aonyx capensis | NA |
|  |  |  |  |  | Civettictis civetta | NA |
|  |  |  |  |  | Crossarchus obscurus | NA |
|  |  |  |  |  | Panthera pardus | NA |
| Mazurek and Zielinski, 2004 | USA | Unspecified | Species presence | Yes | Lynx rufus | NA |
|  |  |  |  |  | Mephitis mephitis | NA |
|  |  |  |  |  | Mustela erminea | NA |
|  |  |  |  |  | Procyon lotor | NA |
|  |  |  |  |  | Spilogale gracilis | NA |
|  |  |  |  |  | Urocyon cinereoargenteus | NA |
|  |  |  |  |  | Ursus americanus | NA |
| McKay and Finnegan, 2022 | Canada | Clearfelling | Occupancy | Yes | Canis lupus | NA |
|  |  |  |  |  | Puma concolor | NA |
|  |  |  |  |  | Ursus americanus | NA |
|  |  |  |  |  | Ursus arctos | Unharvested forest |
| McNitt et al., 2020 | USA | Mixed | Habitat use | Yes | Lynx rufus | NA |
| Mohamad et al., 2015 | Malaysia | Partial harvesting | Density | Yes | Neofelis nebulosa | No difference |
| Mohamed et al., 2013 | Malaysia | Partial harvesting | Density | Yes | Prionailurus bengalensis | NA |
| Mohamed et al., 2021 | Malaysia | Mixed | Density | Yes | Neofelis diardi | No difference |
| Mohd‐Azlan and Sharma, 2003 | Malaysia | Unspecified | Density | Yes | Panthera tigris | NA |
| Mohd-Azlan et al., 2022 | Malaysia | Unspecified | Probability of occurrence | Yes | Catopuma badia | Harvested forest |
|  |  |  |  |  | Neofelis diardi | Unharvested forest |
|  |  |  |  |  | Pardofelis marmorata | Unharvested forest |
|  |  |  |  |  | Prionailurus bengalensis | Harvested forest |
| Mowat and Poole, 2005 | Canada | Clearfelling | Habitat use | Yes | Mustela erminea | NA |
| Muhly et al., 2019 | Canada | Clearfelling | Habitat selection | Yes | Canis lupus | NA |
| Mumma et al., 2018 | Canada | Clearfelling | Habitat use | Yes | Canis lupus | Harvested forest |
| Nakashima et al., 2013 | Malaysia | Unspecified | Habitat use | Yes | Paradoxurus hermaphroditus | NA |
| Nichols, 2017 | USA | Unspecified | Kill site selection | No | Puma concolor | NA |
| Olson et al., 2023 | USA | Mixed | Habitat use | Yes | Lynx canadensis | Harvested forest |
| Organ et al., 2008 | USA | Unspecified | Den site selection | Yes | Lynx canadensis | NA |
| Owen et al., 2015a | USA | Mixed | Habitat use | Yes | Procyon lotor | Unharvested forest |
| Owen et al., 2015b | USA | Mixed | Den site selection | Yes | Procyon lotor | NA |
| Parsons et al., 2020 | USA | Unspecified | Diet | No | Martes pennanti | NA |
| Payer and Harrison, 2003 | USA | Clearfelling | Habitat use | Yes | Martes americana | NA |
| Payer et al., 2005 | USA | Clearfelling | Density | Yes | Martes americana | Unharvested forest |
| Person and Russell, 2009 | USA | Clearfelling | Den site selection | Yes | Canis lupus | Unharvested forest |
| Phoebus et al., 2017 | Canada | Clearfelling | Habitat use | Yes | Ursus arctos | Unharvested forest |
| Pigeon et al., 2016 | Canada | Clearfelling | Habitat use | Yes | Ursus arctos | NA |
| Potvin et al., 1999 | Canada | Clearfelling | Abundance | Yes | Martes americana | Unharvested forest |
| Potvin et al., 2000 | Canada | Clearfelling | Home range | Yes | Martes americana | Unharvested forest |
| Purcell et al., 2009 | USA | Partial harvesting | Den site selection | Yes | Martes pennanti | NA |
| Rajaratnam et al., 2007 | Malaysia | Partial harvesting | Habitat use | Yes | Prionailurus bengalensis | NA |
| Rayan and Linkie, 2015 | Malaysia | Partial harvesting | Density | Yes | Panthera tigris | Unharvested forest |
| Rayan and Linkie, 2016 | Malaysia | Partial harvesting | Interspecific competition | No | Cuon alpinus | NA |
|  |  |  |  |  | Panthera pardus | Harvested forest |
| Rayan and Mohamad, 2009 | Malaysia | Partial harvesting | Density | Yes | Panthera tigris | NA |
| Reagan, 1991 | USA | Clearfelling | Habitat use | Yes | Ursus americanus | NA |
| Roffler et al., 2018 | USA | Clearfelling | Habitat use | Yes | Canis lupus | Unharvested forest |
| Roopsind et al., 2017 | Guyana | Reduced impact logging | Occupancy | Yes | Eira barbara | NA |
|  |  |  |  |  | Leopardus pardalis | No difference |
|  |  |  |  |  | Leopardus wiedii | No difference |
|  |  |  |  |  | Nasua nasua | NA |
|  |  |  |  |  | Panthera onca | No difference |
|  |  |  |  |  | Puma concolor | No difference |
| Roulston, 2013 | USA | Unspecified | Habitat use | Yes | Canis latrans | NA |
|  |  |  |  |  | Canis lupus | NA |
|  |  |  |  |  | Ursus americanus | NA |
| Rozylowicz et al., 2011 | Romania | Clearfelling | Presence/absence | Yes | Felis silvestris | NA |
|  |  |  |  |  | Martes martes | NA |
| Samejima and Semiadi, 2012 | Indonesia | Reduced impact logging | Presence | Yes | Arctictis binturong | NA |
|  |  |  |  |  | Catopuma badia | NA |
|  |  |  |  |  | Cynogale bennettii | NA |
|  |  |  |  |  | Diplogale hosei | NA |
|  |  |  |  |  | Helarctos malayanus | NA |
|  |  |  |  |  | Hemigalus derbyanus | NA |
|  |  |  |  |  | Herpestes brachyurus | NA |
|  |  |  |  |  | Herpestes semitorquatus | NA |
|  |  |  |  |  | Martes flavigula | NA |
|  |  |  |  |  | Neofelis diardi | NA |
|  |  |  |  |  | Paguma larvata | NA |
|  |  |  |  |  | Paradoxurus hermaphroditus | NA |
|  |  |  |  |  | Pardofelis marmorata | NA |
|  |  |  |  |  | Prionailurus bengalensis | NA |
|  |  |  |  |  | Prionailurus planiceps | NA |
|  |  |  |  |  | Prionodon linsang | NA |
|  |  |  |  |  | Viverra tangalunga | NA |
| Samejima et al., 2012 | Malaysia | Reduced impact logging | Mean trapping rate | Yes | Arctictis binturong | No difference |
|  |  |  |  |  | Arctogalidia trivirgata | No difference |
|  |  |  |  |  | Catopuma badia | No difference |
|  |  |  |  |  | Cynogale bennettii | No difference |
|  |  |  |  |  | Helarctos malayanus | No difference |
|  |  |  |  |  | Hemigalus derbyanus | No difference |
|  |  |  |  |  | Herpestes brachyurus | No difference |
|  |  |  |  |  | Herpestes semitorquatus | No difference |
|  |  |  |  |  | Martes flavigula | No difference |
|  |  |  |  |  | Mydaus javanensis | No difference |
|  |  |  |  |  | Neofelis diardi | No difference |
|  |  |  |  |  | Paradoxurus hermaphroditus | No difference |
|  |  |  |  |  | Pardofelis marmorata | No difference |
|  |  |  |  |  | Prionailurus bengalensis | No difference |
|  |  |  |  |  | Prionailurus planiceps | No difference |
|  |  |  |  |  | Prionodon linsang | No difference |
|  |  |  |  |  | Viverra tangalunga | No difference |
| Sanders and Cornman, 2017 | USA | Unspecified | Resting sites | Yes | Martes americana | NA |
| Sauder and Rachlow, 2014 | USA | Clearfelling | Habitat use | Yes | Martes pennanti | Unharvested forest |
| Scrafford et al., 2017 | Canada | Clearfelling | Habitat use | Yes | Gulo gulo | Harvested forest |
| Seip, 2021 | Canada | Clearfelling | Occurrence | Yes | Canis latrans | Harvested forest |
|  |  |  |  |  | Canis lupus | Harvested forest |
|  |  |  |  |  | Lynx canadensis | Harvested forest |
|  |  |  |  |  | Ursus americanus | Harvested forest |
|  |  |  |  |  | Ursus arctos | Harvested forest |
| Shwe et al., 2022 | Myanmar | Partial harvesting | Presence/absence | Yes | Cuon alpinus | No difference |
|  |  |  |  |  | Panthera pardus | Unharvested forest |
|  |  |  |  |  | Panthera tigris | No difference |
| Sidorovich et al., 2010 | Belarus | Clearfelling | Diet | No | Martes martes | NA |
|  |  |  |  |  | Vulpes vulpes | NA |
| Simons‐Legaard et al., 2013 | USA | Mixed | Habitat use | Yes | Lynx canadensis | Unharvested forest |
| Simons‐Legaard et al., 2016 | USA | Clearfelling | Probability of occurrence | Yes | Lynx canadensis | Harvested forest |
| Simons-Legaard et al., 2022 | USA | Unspecified | Habitat use | Yes | Martes americana | Unharvested forest |
| Sivertsen et al., 2016 | Sweden | Clearfelling | Habitat use | Yes | Ursus arctos | Unharvested forest |
| Slauson and Zielinski, 2009 | USA | Unspecified | Den site selection | Yes | Martes americana | NA |
| Smallwood, 1994 | USA | Clearfelling | Density | Yes | Puma concolor | Unharvested forest |
| Smith, 2021 | USA | Partial harvesting | Habitat selection | Yes | Martes pennanti | Unharvested forest |
| Snyder and Bissonette, 1987 | Canada | Clearfelling | Habitat use | Yes | Martes americana | No difference |
| Sollmann et al., 2017 | Malaysia | Mixed | Occupancy | Yes | Aonyx cinereus | NA |
|  |  |  |  |  | Catopuma badia | NA |
|  |  |  |  |  | Cynogale bennettii | NA |
|  |  |  |  |  | Helarctos malayanus | NA |
|  |  |  |  |  | Hemigalus derbyanus | NA |
|  |  |  |  |  | Herpestes brachyurus | NA |
|  |  |  |  |  | Herpestes semitorquatus | NA |
|  |  |  |  |  | Lutrogale perspicillata | NA |
|  |  |  |  |  | Martes flavigula | NA |
|  |  |  |  |  | Mydaus javanensis | NA |
|  |  |  |  |  | Paradoxurus hermaphroditus | NA |
|  |  |  |  |  | Prionailurus bengalensis | Harvested forest |
|  |  |  |  |  | Prionailurus planiceps | NA |
|  |  |  |  |  | Viverra tangalunga | NA |
| Soofi, 2018 | Iran | Unspecified | Occupancy | Yes | Canis lupus | NA |
|  |  |  |  |  | Panthera pardus | NA |
|  |  |  |  |  | Ursus arctos | NA |
| Soutiere, 1979 | USA | Mixed | Density | Yes | Martes americana | NA |
| Squires et al., 2010 | Canada | Mixed | Resource selection | Yes | Lynx canadensis | NA |
| Steventon and Major, 1982 | USA | Mixed | Habitat use | Yes | Martes americana | NA |
| Storch et al., 1990 | Sweden | Clearfelling | Habitat use | Yes | Martes martes | Unharvested forest |
| St-Pierre et al., 2022 | Canada | Clearfelling | Road use | No | Canis lupus | NA |
|  |  |  |  |  | Ursus americanus | NA |
| Suffice et al., 2017 | Canada | Mixed | Habitat use | Yes | Martes americana | Unharvested forest |
|  |  |  |  |  | Martes pennanti | NA |
| Suffice et al., 2020 | Canada | Mixed | Abundance | Yes | Martes americana | No difference |
|  |  |  |  |  | Martes pennanti | NA |
| Sweitzer et al., 2016 | USA | Unspecified | Occupancy/persistence | Yes | Martes pennanti | No difference |
| Thompson, 1994 | Canada | Clearfelling | Demography | No | Martes americana | Unharvested forest |
| Tobler et al., 2018 | Guatemala and Peru | Reduced impact logging | Density | Yes | Canis latrans | No difference |
|  |  |  |  |  | Conepatus semistriatus | No difference |
|  |  |  |  |  | Eira barbara | No difference |
|  |  |  |  |  | Herpailurus yagouaroundi | No difference |
|  |  |  |  |  | Leopardus pardalis | No difference |
|  |  |  |  |  | Leopardus wiedii | No difference |
|  |  |  |  |  | Nasua narica | No difference |
|  |  |  |  |  | Panthera onca | No difference |
|  |  |  |  |  | Procyon lotor | No difference |
|  |  |  |  |  | Urocyon cinereoargenteus | No difference |
| Toews et al., 2018 | Canada | Clearfelling | Abundance | Yes | Canis latrans | No difference |
|  |  |  |  |  | Canis lupus | No difference |
|  |  |  |  |  | Lynx canadensis | No difference |
| Tseganu, 2015 | Ghana | Unspecified | Presence/absence | Yes | Crossarchus obscurus | NA |
|  |  |  |  |  | Genetta pardina | NA |
| Twining et al., 2017 | Malaysia | Partial harvesting | Occupancy | Yes | Viverra tangalunga | NA |
| Van Manen and Pelton, 1997 | USA | Unspecified | Habitat use | Yes | Ursus americanus | NA |
| Vanlandeghem et al., 2021 | Canada | Clearfelling | Habitat use | Yes | Canis lupus | Unharvested forest |
| Vashon et al., 2008 | USA | Clearfelling | Habitat use | Yes | Lynx canadensis | NA |
| Wall et al., 2021 | Indonesia | Partial harvesting | Presence/absence | Yes | Aonyx cinereus | NA |
|  |  |  |  |  | Arctictis binturong | NA |
|  |  |  |  |  | Catopuma badia | NA |
|  |  |  |  |  | Cynogale bennettii | NA |
|  |  |  |  |  | Diplogale hosei | NA |
|  |  |  |  |  | Helarctos malayanus | NA |
|  |  |  |  |  | Hemigalus derbyanus | NA |
|  |  |  |  |  | Herpestes brachyurus | NA |
|  |  |  |  |  | Herpestes semitorquatus | NA |
|  |  |  |  |  | Martes flavigula | NA |
|  |  |  |  |  | Neofelis diardi | NA |
|  |  |  |  |  | Paguma larvata | NA |
|  |  |  |  |  | Paradoxurus hermaphroditus | NA |
|  |  |  |  |  | Pardofelis marmorata | NA |
|  |  |  |  |  | Prionailurus bengalensis | NA |
|  |  |  |  |  | Prionodon linsang | NA |
|  |  |  |  |  | Viverra tangalunga | NA |
| Wallgren et al., 2009 | Sweden | Clearfelling | Abundance | Yes | Meles meles | Harvested forest |
| Wampler et al., 2008 | USA | Mixed | Abundance | Yes | Lynx rufus | NA |
|  |  |  |  |  | Mephitis mephitis | NA |
|  |  |  |  |  | Ursus americanus | NA |
| Wasserman et al., 2012 | USA | Clearfelling | Habitat use | Yes | Martes americana | Unharvested forest |
| Watine et al., 2022 | Belize | Partial harvesting | Abundance | Yes | Leopardus pardalis | NA |
|  |  |  |  |  | Panthera onca | NA |
|  |  |  |  |  | Puma concolor | NA |
|  |  |  |  |  | Urocyon cinereoargenteus | NA |
| Wayne et al., 2011 | Australia | Mixed | Presence/absence | Yes | Dasyurus geoffroii | NA |
| Wearn et al., 2013 | Malaysia | Clearfelling | Abundance | Yes | Catopuma badia | NA |
|  |  |  |  |  | Neofelis diardi | NA |
|  |  |  |  |  | Pardofelis marmorata | NA |
|  |  |  |  |  | Prionailurus bengalensis | NA |
|  |  |  |  |  | Prionailurus planiceps | NA |
| Wearn et al., 2016 | Malaysia | Unspecified | Presence/absence | Yes | Arctictis binturong | NA |
|  |  |  |  |  | Catopuma badia | NA |
|  |  |  |  |  | Helarctos malayanus | NA |
|  |  |  |  |  | Hemigalus derbyanus | NA |
|  |  |  |  |  | Herpestes brachyurus | NA |
|  |  |  |  |  | Herpestes semitorquatus | NA |
|  |  |  |  |  | Martes flavigula | NA |
|  |  |  |  |  | Mydaus javanensis | NA |
|  |  |  |  |  | Neofelis diardi | NA |
|  |  |  |  |  | Paguma larvata | NA |
|  |  |  |  |  | Paradoxurus hermaphroditus | NA |
|  |  |  |  |  | Pardofelis marmorata | NA |
|  |  |  |  |  | Prionailurus bengalensis | NA |
|  |  |  |  |  | Prionodon linsang | NA |
|  |  |  |  |  | Viverra tangalunga | NA |
| Wearn et al., 2022 | Malaysia | Unspecified | Density and activity | Yes | Aonyx cinereus | No difference |
|  |  |  |  |  | Arctictis binturong | Unharvested forest |
|  |  |  |  |  | Catopuma badia | Harvested forest |
|  |  |  |  |  | Diplogale hosei | Unharvested forest |
|  |  |  |  |  | Helarctos malayanus | Harvested forest |
|  |  |  |  |  | Hemigalus derbyanus | NA |
|  |  |  |  |  | Herpestes brachyurus | No difference |
|  |  |  |  |  | Herpestes semitorquatus | No difference |
|  |  |  |  |  | Martes flavigula | No difference |
|  |  |  |  |  | Mustela nudipes | No difference |
|  |  |  |  |  | Mydaus javanensis | Harvested forest |
|  |  |  |  |  | Neofelis diardi | No difference |
|  |  |  |  |  | Paguma larvata | Unharvested forest |
|  |  |  |  |  | Pardofelis marmorata | No difference |
|  |  |  |  |  | Prionailurus bengalensis | No difference |
|  |  |  |  |  | Viverra tangalunga | NA |
| Wegge et al., 2019 | Norway | Clearfelling | Density | Yes | Vulpes vulpes | NA |
| Wei et al., 2015 | Malaysia | Partial harvesting | Presence | Yes | Panthera pardus | NA |
| Wening et al., 2019 | Germany | Clearfelling | Detection | Yes | Felis silvestris | NA |
| White et al., 2001 | USA | Unspecified | Den use | Yes | Ursus americanus | NA |
| Wibisono et al., 2018 | Indonesia | Unspecified | Occurrence | Yes | Panthera pardus | Unharvested forest |
| Willcox et al., 2012 | Vietnam | Unspecified | Presence | Yes | Arctogalidia trivirgata | NA |
| Wilting et al., 2010 | Malaysia | Reduced impact logging | Presence | Yes | Aonyx cinereus | NA |
|  |  |  |  |  | Arctictis binturong | NA |
|  |  |  |  |  | Arctogalidia trivirgata | NA |
|  |  |  |  |  | Cynogale bennettii | NA |
|  |  |  |  |  | Hemigalus derbyanus | NA |
|  |  |  |  |  | Herpestes brachyurus | NA |
|  |  |  |  |  | Herpestes semitorquatus | NA |
|  |  |  |  |  | Lutra sumatrana | NA |
|  |  |  |  |  | Lutrogale perspicillata | NA |
|  |  |  |  |  | Martes flavigula | NA |
|  |  |  |  |  | Mydaus javanensis | NA |
|  |  |  |  |  | Paradoxurus hermaphroditus | NA |
|  |  |  |  |  | Prionodon linsang | NA |
|  |  |  |  |  | Viverra tangalunga | NA |
| Wilting et al., 2012 | Malaysia | Reduced impact logging | Density | Yes | Neofelis diardi | NA |
| Wong and Linkie, 2013 | Indonesia | Partial harvesting | Occupancy | Yes | Helarctos malayanus | Unharvested forest |
| Wong et al., 2022 | Malaysia | Mixed | Occupancy | Yes | Aonyx cinereus | NA |
|  |  |  |  |  | Catopuma badia | NA |
|  |  |  |  |  | Diplogale hosei | NA |
|  |  |  |  |  | Helarctos malayanus | NA |
|  |  |  |  |  | Hemigalus derbyanus | NA |
|  |  |  |  |  | Martes flavigula | NA |
|  |  |  |  |  | Paguma larvata | NA |
|  |  |  |  |  | Paradoxurus hermaphroditus | NA |
|  |  |  |  |  | Pardofelis marmorata | NA |
|  |  |  |  |  | Prionailurus bengalensis | NA |
|  |  |  |  |  | Viverra tangalunga | NA |
| Yi et al., 2022 | Malaysia | Unspecified | Occurrence | Yes | Arctictis binturong | No difference |
|  |  |  |  |  | Arctogalidia trivirgata | No difference |
|  |  |  |  |  | Helarctos malayanus | Harvested forest |
|  |  |  |  |  | Hemigalus derbyanus | No difference |
|  |  |  |  |  | Herpestes brachyurus | No difference |
|  |  |  |  |  | Martes flavigula | No difference |
|  |  |  |  |  | Prionailurus bengalensis | Unharvested forest |
|  |  |  |  |  | Viverra tangalunga | No difference |
| Zamuda et al., 2022 | USA | Partial harvesting | Occupancy | Yes | Canis latrans | No difference |
|  |  |  |  |  | Lynx rufus | Harvested forest |
|  |  |  |  |  | Martes pennanti | Unharvested forest |
|  |  |  |  |  | Procyon lotor | No difference |
| Zhang et al., 2019 | China | Unspecified | Habitat use | Yes | Arctonyx albogularis | Unharvested forest |
|  |  |  |  |  | Martes flavigula | Unharvested forest |
|  |  |  |  |  | Mustela sibirica | Unharvested forest |
|  |  |  |  |  | Paguma larvata | Unharvested forest |
|  |  |  |  |  | Prionailurus bengalensis | NA |
|  |  |  |  |  | Ursus thibetanus | Unharvested forest |
| Zhou et al., 2008 | China | Mixed | Diet | No | Melogale moschata | NA |
| Zhou et al., 2015 | China | Mixed | Latrine use | No | Arctonyx collaris | Harvested forest |
| Zielinski et al., 2013 | USA | Mixed | Abundance | Yes | Martes pennanti | NA |

**Appendix S3:** List of native mammalian carnivore species recorded in production forests, along with species traits. Body size and diet for each species were identified using the Encyclopedia of Life (https://eol.org/) and Animal Diversity Web (https://animaldiversity.org/).

| Species name | Common name | Family | Recorded in plantation? | Recorded in harvested native forest? | IUCN Red List Category | Body size (kg) | Diet |
| --- | --- | --- | --- | --- | --- | --- | --- |
| Aonyx capensis | Cape clawless otter | Mustelidae | Yes | Yes | Near Threatened | 19 | Piscivore |
| Aonyx cinereus | Small-clawed otter | Mustelidae | Yes | Yes | Vulnerable | 1 | Piscivore |
| Arctictis binturong | Binturong | Viverridae | No | Yes | Vulnerable | 15 | Omnivore |
| Arctogalidia trivirgata | Small-toothed palm civet | Viverridae | Yes | Yes | Least Concern | 2.25 | Omnivore |
| Arctonyx albogularis | Northern hog badger | Mustelidae | No | Yes | Least Concern | 6.36 | Omnivore |
| Arctonyx collaris | Hog badger | Mustelidae | No | Yes | Vulnerable | 6.36 | Omnivore |
| Atelocynus microtis | Short-eared dog | Canidae | No | Yes | Near Threatened | 9.5 | Omnivore |
| Atilax paludinosus | Water mongoose | Herpestidae | Yes | Yes | Least Concern | 3.3 | Omnivore |
| Bdeogale nigripes | Black-legged mongoose | Herpestidae | No | Yes | Least Concern | 2.5 | Insectivore |
| Canis adustus | Side-striped jackal | Canidae | Yes | No | Least Concern | 10.25 | Omnivore |
| Canis aureus | Golden jackal | Canidae | Yes | No | Least Concern | 9 | Omnivore |
| Canis latrans | Coyote | Canidae | No | Yes | Least Concern | 11.9 | Omnivore |
| Canis lupus | Grey wolf | Canidae | Yes | Yes | Least Concern | 30.75 | Hypercarnivore |
| Canis mesomelas | Black-backed jackal | Canidae | Yes | No | Least Concern | 8.5 | Omnivore |
| Caracal aurata | African golden cat | Felidae | No | Yes | Vulnerable | 11 | Hypercarnivore |
| Caracal caracal | Caracal | Felidae | Yes | No | Least Concern | 13.75 | Hypercarnivore |
| Catopuma badia | Bay cat | Felidae | Yes | Yes | Endangered | 4 | Hypercarnivore |
| Catopuma temminckii | Asian golden cat | Felidae | No | Yes | Near Threatened | 11.5 | Hypercarnivore |
| Cerdocyon thous | Crab-eating fox | Canidae | Yes | No | Least Concern | 8.2 | Omnivore |
| Chrysocyon brachyurus | Maned wolf | Canidae | Yes | No | Near Threatened | 24 | Omnivore |
| Civettictis civetta | African civet | Viverridae | Yes | Yes | Least Concern | 12 | Omnivore |
| Conepatus chinga | Molina's hog-nosed skunk | Mephitidae | Yes | No | Least Concern | 1.92 | Omnivore |
| Conepatus semistriatus | Striped hog-nosed skunk | Mephitidae | Yes | Yes | Least Concern | 1.2 | Omnivore |
| Crocuta crocuta | Spotted hyena | Hyaenidae | Yes | No | Least Concern | 63 | Hypercarnivore |
| Crossarchus obscurus | Common kusimanse | Herpestidae | No | Yes | Least Concern | 1.25 | Insectivore |
| Crossarchus platycephalus | Flat-headed cusimanse | Herpestidae | No | Yes | Least Concern | 1 | Omnivore |
| Cryptoprocta ferox | Fossa | Eupleridae | No | Yes | Vulnerable | 9.5 | Hypercarnivore |
| Cuon alpinus | Dhole | Canidae | Yes | Yes | Endangered | 12.76 | Omnivore |
| Cynogale bennettii | Otter civet | Viverridae | No | Yes | Endangered | 4.5 | Omnivore |
| Dasyurus geoffroii | Western quoll | Dasyuridae | No | Yes | Near Threatened | 1.1 | Omnivore |
| Dasyurus maculatus | Spotted-tailed quoll | Dasyuridae | Yes | Yes | Near Threatened | 5.5 | Hypercarnivore |
| Dasyurus viverrinus | Eastern quoll | Dasyuridae | Yes | Yes | Endangered | 1.09 | Omnivore |
| Diplogale hosei | Hose's palm civet | Viverridae | Yes | Yes | Vulnerable | 5.45 | Omnivore |
| Eira barbara | Tayra | Mustelidae | Yes | Yes | Least Concern | 5.05 | Omnivore |
| Eupleres goudotii | Small-toothed civet | Eupleridae | No | Yes | Vulnerable | 3 | Insectivore |
| Felis chaus | Jungle cat | Felidae | Yes | No | Least Concern | 7 | Hypercarnivore |
| Felis silvestris | European wildcat | Felidae | Yes | Yes | Least Concern | 4.25 | Hypercarnivore |
| Fossa fossana | Malagasy civet | Eupleridae | No | Yes | Vulnerable | 1.5 | Omnivore |
| Galerella sanguineus | Slender mongoose | Herpestidae | Yes | No | Least Concern | 0.55 | Omnivore |
| Galictis cuja | Lesser grison | Mustelidae | Yes | No | Least Concern | 1.36 | Omnivore |
| Galidia elegans | Ring-tailed mongoose | Eupleridae | No | Yes | Least Concern | 0.8 | Omnivore |
| Galidictis fasciata | Broad-striped mongoose | Eupleridae | No | Yes | Vulnerable | 0.55 | Hypercarnivore |
| Genetta genetta | Common genet | Viverridae | Yes | No | Least Concern | 1.6 | Hypercarnivore |
| Genetta pardina | Pardine genet | Viverridae | No | Yes | Least Concern | 2 | Omnivore |
| Genetta servalina | Servaline genet | Viverridae | No | Yes | Least Concern | 1 | Omnivore |
| Genetta tigrina | Large spotted genet | Viverridae | Yes | No | Least Concern | 2.23 | Omnivore |
| Gulo gulo | Wolverine | Mustelidae | No | Yes | Least Concern | 14.53 | Hypercarnivore |
| Helarctos malayanus | Sun bear | Ursidae | Yes | Yes | Vulnerable | 46 | Omnivore |
| Hemigalus derbyanus | Banded palm civet | Viverridae | Yes | Yes | Near Threatened | 2.5 | Omnivore |
| Herpailurus yagouaroundi | Jaguarundi | Felidae | Yes | Yes | Least Concern | 6.3 | Hypercarnivore |
| Herpestes brachyurus | Short-tailed mongoose | Herpestidae | Yes | Yes | Near Threatened | 1.41 | Omnivore |
| Herpestes ichneumon | Egyptian mongoose | Herpestidae | Yes | No | Least Concern | 2.85 | Omnivore |
| Herpestes naso | Long-nosed mongoose | Herpestidae | No | Yes | Least Concern | 3 | Omnivore |
| Herpestes semitorquatus | Collared mongoose | Herpestidae | Yes | Yes | Near Threatened | 1.5 | Omnivore |
| Herpestes urva | Crab-eating mongoose | Herpestidae | No | Yes | Least Concern | 2 | Omnivore |
| Ichneumia albicauda | White-tailed mongoose | Herpestidae | Yes | No | Least Concern | 3.5 | Insectivore |
| Ictonyx striatus | Striped polecat | Mustelidae | Yes | No | Least Concern | 1.3 | Hypercarnivore |
| Leopardus colocolo | Colo colo | Felidae | Yes | No | Near Threatened | 4 | Hypercarnivore |
| Leopardus geoffroyi | Geoffroy's cat | Felidae | Yes | No | Least Concern | 3.59 | Hypercarnivore |
| Leopardus guigna | Kodkod | Felidae | Yes | No | Vulnerable | 2.23 | Hypercarnivore |
| Leopardus guttulus | Southern tiger cat | Felidae | Yes | No | Vulnerable | 2.5 | Hypercarnivore |
| Leopardus pardalis | Ocelot | Felidae | Yes | Yes | Least Concern | 6.3 | Hypercarnivore |
| Leopardus tigrinus | Oncilla | Felidae | Yes | No | Vulnerable | 2.25 | Hypercarnivore |
| Leopardus wiedii | Margay | Felidae | Yes | Yes | Near Threatened | 2.38 | Hypercarnivore |
| Leptailurus serval | Serval | Felidae | Yes | No | Least Concern | 12 | Hypercarnivore |
| Lontra longicaudis | Neotropical otter | Mustelidae | Yes | No | Near Threatened | 8.1 | Piscivore |
| Lutra lutra | Eurasian otter | Mustelidae | Yes | No | Near Threatened | 11 | Piscivore |
| Lutra sumatrana | Hairy-nosed otter | Mustelidae | No | Yes | Endangered | 5.5 | Piscivore |
| Lutrogale perspicillata | Smooth-coated otter | Mustelidae | No | Yes | Vulnerable | 9 | Piscivore |
| Lycalopex culpaeus | Culpeo | Canidae | Yes | No | Least Concern | 9.8 | Omnivore |
| Lycalopex fulvipes | Darwin's fox | Canidae | Yes | No | Endangered | 2.7 | Omnivore |
| Lycalopex griseus | South American grey fox | Canidae | Yes | No | Least Concern | 3 | Omnivore |
| Lycalopex gymnocercus | Pampas fox | Canidae | Yes | No | Least Concern | 4 | Omnivore |
| Lycaon pictus | Painted wolf | Canidae | Yes | No | Endangered | 26.8 | Hypercarnivore |
| Lynx canadensis | Canada lynx | Felidae | No | Yes | Least Concern | 9.37 | Hypercarnivore |
| Lynx lynx | Lynx | Felidae | Yes | Yes | Least Concern | 17.95 | Hypercarnivore |
| Lynx pardinus | Iberian lynx | Felidae | Yes | No | Endangered | 9.4 | Hypercarnivore |
| Lynx rufus | Bobcat | Felidae | No | Yes | Least Concern | 8.9 | Hypercarnivore |
| Martes americana | American marten | Mustelidae | No | Yes | Least Concern | 1.25 | Hypercarnivore |
| Martes flavigula | Yellow-throated marten | Mustelidae | Yes | Yes | Least Concern | 2.5 | Omnivore |
| Martes foina | Stone marten | Mustelidae | Yes | Yes | Least Concern | 1.54 | Omnivore |
| Martes martes | Pine marten | Mustelidae | Yes | Yes | Least Concern | 1.3 | Omnivore |
| Martes melampus | Japanese marten | Mustelidae | Yes | No | Least Concern | 1 | Omnivore |
| Martes pennanti | Fisher | Mustelidae | No | Yes | Least Concern | 4 | Hypercarnivore |
| Martes zibellina | Sable | Mustelidae | No | Yes | Least Concern | 1.13 | Omnivore |
| Meles anakuma | Japanese badger | Mustelidae | Yes | No | Least Concern | 5.8 | Insectivore |
| Meles meles | European badger | Mustelidae | Yes | Yes | Least Concern | 13 | Omnivore |
| Mellivora capensis | Honey badger | Mustelidae | Yes | No | Least Concern | 9 | Omnivore |
| Melogale moschata | Chinese ferret-badger | Mustelidae | No | Yes | Least Concern | 2 | Omnivore |
| Melursus ursinus | Sloth bear | Ursidae | Yes | No | Vulnerable | 100 | Omnivore |
| Mephitis mephitis | Striped skunk | Mephitidae | No | Yes | Least Concern | 2.09 | Omnivore |
| Mungos mungo | Banded mongoose | Herpestidae | Yes | No | Least Concern | 1.9 | Insectivore |
| Mustela erminea | Stoat | Mustelidae | Yes | Yes | Least Concern | 0.17 | Hypercarnivore |
| Mustela frenata | Long-tailed weasel | Mustelidae | No | Yes | Least Concern | 0.15 | Hypercarnivore |
| Mustela itatsi | Japanese weasel | Mustelidae | Yes | No | Near Threatened | 0.4 | Hypercarnivore |
| Mustela lutreola | European mink | Mustelidae | Yes | No | Critically Endangered | 0.44 | Hypercarnivore |
| Mustela nivalis | Weasel | Mustelidae | Yes | Yes | Least Concern | 0.1 | Hypercarnivore |
| Mustela nudipes | Malay weasel | Mustelidae | Yes | Yes | Least Concern | 0.5 | Hypercarnivore |
| Mustela putorius | Polecat | Mustelidae | Yes | No | Least Concern | 0.73 | Hypercarnivore |
| Mustela sibirica | Siberian weasel | Mustelidae | Yes | Yes | Least Concern | 0.41 | Omnivore |
| Mydaus javanensis | Malay badger | Mephitidae | Yes | Yes | Least Concern | 2.5 | Insectivore |
| Nandinia binotata | African civet | Nandiniidae | No | Yes | Least Concern | 2 | Omnivore |
| Nasua narica | White-nosed coati | Procyonidae | Yes | Yes | Least Concern | 4.03 | Omnivore |
| Nasua nasua | South American coati | Procyonidae | Yes | Yes | Least Concern | 3.3 | Omnivore |
| Neofelis diardi | Sunda clouded leopard | Felidae | No | Yes | Vulnerable | 20 | Hypercarnivore |
| Neofelis nebulosa | Clouded leopard | Felidae | No | Yes | Vulnerable | 19.5 | Hypercarnivore |
| Nyctereutes procyonoides | Raccoon dog | Canidae | Yes | No | Least Concern | 4.04 | Omnivore |
| Paguma larvata | Masked palm civet | Viverridae | Yes | Yes | Least Concern | 4.3 | Omnivore |
| Panthera onca | Jaguar | Felidae | Yes | Yes | Near Threatened | 94.5 | Hypercarnivore |
| Panthera pardus | Leopard | Felidae | Yes | Yes | Vulnerable | 55.62 | Hypercarnivore |
| Panthera tigris | Tiger | Felidae | Yes | Yes | Endangered | 162.56 | Hypercarnivore |
| Paradoxurus hermaphroditus | Asian palm civet | Viverridae | Yes | Yes | Least Concern | 3.2 | Omnivore |
| Pardofelis marmorata | Marbled cat | Felidae | Yes | Yes | Near Threatened | 3.25 | Hypercarnivore |
| Prionailurus bengalensis | Leopard cat | Felidae | Yes | Yes | Least Concern | 3.3 | Hypercarnivore |
| Prionailurus planiceps | Flat-headed cat | Felidae | No | Yes | Endangered | 6.75 | Hypercarnivore |
| Prionodon linsang | Banded linsang | Prionodontidae | No | Yes | Least Concern | 0.7 | Omnivore |
| Procyon cancrivorus | Crab-eating raccoon | Procyonidae | Yes | Yes | Least Concern | 7.8 | Omnivore |
| Procyon lotor | Raccoon | Procyonidae | Yes | Yes | Least Concern | 5.53 | Omnivore |
| Puma concolor | Puma | Felidae | Yes | Yes | Least Concern | 33 | Hypercarnivore |
| Rhynchogale melleri | Meller's mongoose | Herpestidae | Yes | No | Least Concern | 2.5 | Insectivore |
| Sarcophilus harrisii | Tasmanian devil | Dasyuridae | Yes | Yes | Endangered | 8 | Hypercarnivore |
| Speothos venaticus | Bush dog | Canidae | No | Yes | Near Threatened | 6 | Hypercarnivore |
| Spilogale gracilis | Western spotted skunk | Mephitidae | No | Yes | Least Concern | 4.6 | Omnivore |
| Spilogale putorius | Allegheny spotted skunk | Mephitidae | No | Yes | Vulnerable | 0.34 | Omnivore |
| Urocyon cinereoargenteus | Grey fox | Canidae | No | Yes | Least Concern | 3.8 | Omnivore |
| Ursus americanus | Black bear | Ursidae | Yes | Yes | Least Concern | 100 | Omnivore |
| Ursus arctos | Brown bear | Ursidae | Yes | Yes | Least Concern | 206 | Omnivore |
| Ursus thibetanus | Asiatic black bear | Ursidae | Yes | Yes | Vulnerable | 77.5 | Omnivore |
| Viverra tangalunga | Malay civet | Viverridae | Yes | Yes | Least Concern | 10 | Omnivore |
| Viverricula indica | Small Indian civet | Viverridae | Yes | No | Least Concern | 2.98 | Omnivore |
| Vulpes bengalensis | Indian fox | Canidae | Yes | No | Least Concern | 2.73 | Omnivore |
| Vulpes vulpes | Red fox | Canidae | Yes | Yes | Least Concern | 5.95 | Omnivore |

**Appendix S4:** Responses of carnivore species to habitat features in production forests, including the studies recording these results.

| **Habitat feature** | **Plantations** | | **Harvested native forest** | |
| --- | --- | --- | --- | --- |
|  | *Positive response* | *Negative response* | *Positive response* | *Negative response* |
| Stand age |  |  |  |  |
| - Younger |  |  | Crimmins et al., 2012  Fuller and Harrison, 2005  Krebs et al., 2007  Lesmeister et al., 2013  Lisgo, 1999  Lomolino and Perault, 2000  Mathai et al., 2017  McNitt et al., 2020  Mohamed et al., 2013  Mowat and Poole, 2005  Roffler et al., 2018  Sivertsen et al., 2016 |  |
| - Mid-stage regenerating |  |  | Belcher, 2008  Boisjoly et al., 2010  Hearn et al., 2010  Hoving et al., 2004  Lomolino and Perault  2000  Olson et al., 2023  Payer and Harrison  2003 |  |
| - Older/mature |  |  | Augeri, 2005  Bull et al., 2005  Buskirk et al., 1996  Holbrook et al., 2017, 2018, 2019  Kosterman et al., 2018  Owen et al., 2015a  Sauder and Rachlow, 2014  Simons‐Legaard et al., 2013  Soutiere, 1979  Steventon and Major, 1982  Storch et al., 1990  Vanlandeghem et al., 2021  Wasserman et al., 2012 |  |
| Clearcuts | Brandt and Lambin, 2007  Eom et al., 2019  Escudero-Páez et al., 2018  Linnell et al., 2017 |  | Boisjoly et al., 2010  Crimmins et al., 2012  Fuller and Harrison, 2005  Kertson and Marzluff, 2011  Lisgo, 1999 | Fuller and Harrison, 2010  Happe et al., 2020  Hoving et al., 2004  Lesmerises et al., 2012  Potvin et al., 2000  Sivertsen et al., 2016  Snyder and Bissonette, 1987  Soutiere, 1979  Storch et al., 1990 |
| Undergrowth | Acosta-Jamett and Simonetti, 2004  Bojarska et al., 2021  Escudero-Páez et al., 2018  Karelus et al., 2018  Lantschner et al., 2012  Llaneza et al., 2016  Lyall, 2017  Moreira-Arce et al., 2016  Rosalino et al., 2004  Santos and Beier, 2008  Simonetti et al., 2013  Stratman et al., 2001  Sunarto et al., 2012  Takahata et al., 2013 | Jones et al., 2023  Moreira-Arce et al., 2016  Tomita and Hiura, 2021 | Lesmeister et al., 2013  Mathai et al., 2017  Organ et al., 2008  Potvin et al., 2000, 1999  Squires et al., 2010 | Jones et al., 2023  Kunkel and Pletscher, 2000  Payer and Harrison, 2003  Roffler et al., 2018  Watine et al., 2022 |
| Canopy cover | Karelus et al., 2018  Llaneza et al., 2016  Rhim et al., 2015 | Caryl et al., 2012  Linnell et al., 2017  Stratman et al., 2001 | Augeri, 2005  Bull et al., 2005  Buskirk et al., 1996  Fuller and Harrison, 2005  Kordosky et al., 2021  Mathai et al., 2017  Payer and Harrison, 2003  Person and Russell, 2009  Purcell et al., 2009  Sanders and Cornman, 2017  Smith, 2021  Sweitzer et al., 2016  Wasserman et al., 2012  Zamuda et al., 2022 | Kays et al., 2008  McNitt et al., 2020  Mohamed et al., 2013  Mowat and Poole, 2005  Watine et al., 2022  Zamuda et al., 2022 |
| Coarse woody debris | Hwang et al., 2014  Jones et al., 2023  Lyall, 2017  Rhim et al., 2015  Santos and Beier, 2008 |  | Bull et al., 2005  Buskirk et al., 1996  Lisgo, 1999  Lomolino and Perault, 2000  Organ et al., 2008  Payer and Harrison, 2003  Suffice et al., 2017 |  |
| Larger/taller trees | Lyall, 2017  Moreira-Arce et al., 2016  Yamada and Fujioka, 2010 | Calkoen et al., 2018  Moreira-Arce et al., 2016 | Augeri, 2005  Hearn et al., 2010  Mathai et al., 2017  Owen et al., 2015b  Payer and Harrison, 2003  Sanders and Cornman, 2017 |  |
| Riparian areas | Fournier et al., 2007  Karelus et al., 2018  Katna et al., 2022  Ramesh et al., 2016  Rosalino et al., 2004  Santos & Beier, 2008  Takahata et al., 2013  Zabala et al., 2005 |  | Bobo et al., 2017  Bojarska et al., 2017  Bull et al., 2005  Dijak & Thompson, 2000  Gurarie et al., 2011  Homkes, 2021  Kays et al., 2008  Kuzyk et al., 2004  Mohd-Azlan et al., 2022  Owen, Berl, & Edwards, 2015  Person & Russell, 2009  Phoebus et al., 2017  Purcell et al., 2009  Sollmann et al., 2017  Zamuda et al., 2022 |  |
| Forestry roads | Acosta-Jamett and Simonetti, 2004  Paolino et al., 2018  Sunarto et al., 2012 | Acosta-Jamett and Simonetti, 2004  Bojarska et al., 2021  Karelus et al., 2018  Llaneza et al., 2016 | Bowman et al., 2010  Gurarie et al., 2011  Krebs et al., 2007  Lesmerises et al., 2012  Mathai et al., 2017  Mohamed et al., 2013  Mumma et al., 2018  Nichols, 2017  Reagan, 1991  Seip, 2021  Sollmann et al., 2017  Vanlandeghem et al., 2021  Wearn et al., 2013  Zamuda et al., 2022 | Bourbonnais, 2013  Bowman et al., 2010  Kays et al., 2008  Kortello et al., 2019  Kunkel and Pletscher, 2000  Linkie et al., 2008  Mathai et al., 2017  McKay and Finnegan, 2022  Mohd-Azlan et al., 2022  Person and Russell, 2009  Sivertsen et al., 2016  Sollmann et al., 2017  Wong and Linkie, 2013  Zamuda et al., 2022 |
| Habitat edges | Lyra-Jorge et al., 2010  Pedersen et al., 2010 | Sunarto et al., 2012 | Bojarska et al., 2017  Brodie et al., 2015  Gurarie et al., 2011  Houle et al., 2010  Kays et al., 2008  McNitt et al., 2020  Person and Russell, 2009  Scrafford et al., 2017  Simons‐Legaard et al., 2013 | Augeri, 2005  Gulsby et al., 2017  Hargis et al., 1999  Potvin et al., 2000  Wibisono et al., 2018 |
| Den features |  |  |  |  |
| - Large trees |  |  | Buskirk et al., 1996  Owen et al., 2015b  Purcell et al., 2009  Sanders and Cornman, 2017 |  |
| - Logs |  |  | Bull et al., 2005  Bull and Heater, 2000  Buskirk et al., 1996  Slauson and Zielinski, 2009  Steventon and Major, 1982 |  |
| - Tree cavities |  |  | Bull et al., 2005  Bull and Heater, 2000  Buskirk et al., 1996  Nakashima et al., 2013  Sanders and Cornman, 2017 |  |
| - Dead trees/stumps |  |  | Purcell et al., 2009  Slauson and Zielinski, 2009  Steventon and Major, 1982 |  |
| - Logging debris |  |  | Bull and Heater, 2000  Jokinen et al., 2019  Lisgo, 1999  Owen et al., 2015b  Reagan, 1991  Scrafford et al., 2017  Slauson and Zielinski, 2009  White et al., 2001 |  |

**Appendix S5:** Studies recording other results reported in the review.

| Reported result | References |
| --- | --- |
| Carnivore use of other land uses compared to harvested native forest |  |
| - Agricultural land (including oil palm) |  |
| - Positive response | Dijak and Thompson, 2000  Rajaratnam et al., 2007  Toews et al., 2018 |
| - Negative response | Augeri, 2005  Hearn et al., 2019, 2016  Imron et al., 2011  Kays et al., 2008  Toews et al., 2018  Zhou et al., 2015 |
| - Human settlements |  |
| - Positive response | Mathai et al., 2017 |
| - Negative response | Ausilio et al., 2022  Imron et al., 2011  Kaartinen et al., 2015  Kordosky et al., 2021  Lesmerises et al., 2012  Mathai et al., 2017  Shwe et al., 2022  Van Manen and Pelton, 1997 |
| No response to roads by carnivores in production forests | Carvalho Jr et al., 2021  Squires et al., 2010  Tobler et al., 2018  Wearn et al., 2013  Wening et al., 2019  Yi et al., 2022 |
| Studies considering reproductive success of carnivores in production forest | Belcher, 2003  Flynn et al., 2011  Hearn et al., 2016  Holbrook et al., 2019  Kosterman et al., 2018  Vashon et al., 2008  White et al., 2001 |
